# An Intermediate Mesoderm Premyogenic Niche Supports Early Human Myogenic Lineage Progression

**DOI:** 10.64898/2026.03.28.715044

**Authors:** Olga G. Jaime, Katherine F. Bazan, Angela Li, Asja A. Deai, Anita Lakatos, Michael R. Hicks

## Abstract

Transient cell states that precede and support human myogenic lineage commitment, and the intrinsic and extrinsic signals that control them, remain poorly defined in vitro. Here, we used longitudinal single-nucleus profiling, together with a SIX1:H2B-GFP hPSC reporter for lineage tracing, resolved previously uncaptured transient intermediates and sequential waves of human myogenesis across differentiation and *in vivo*. We show that hPSC-directed myogenesis gives rise in parallel to paraxial mesoderm and a transient PAX8+ intermediate mesoderm population that forms a 3-dimensional pre-myogenic niche supporting the PAX3-to-PAX7 myogenic progenitor transition. LIANA+ analysis further identified a temporally restricted BMP7-BMPR1B interaction, together with laminin-linked signaling, between PAX8+ niche cells and skeletal muscle progenitors before commitment. We further show that dynamic SIX1 cofactor switching, including EYA3 activity, is required for PAX3-to-PAX7 progression, and that disruption of this program compromises multi-lineage niche integrity. Together, these findings define how transient niche populations and intrinsic regulatory networks coordinate early human myogenic lineage progression and provide a human in vitro platform to study parallel intermediate and paraxial mesoderm development during myogenesis.

**Graphical Abstract:** 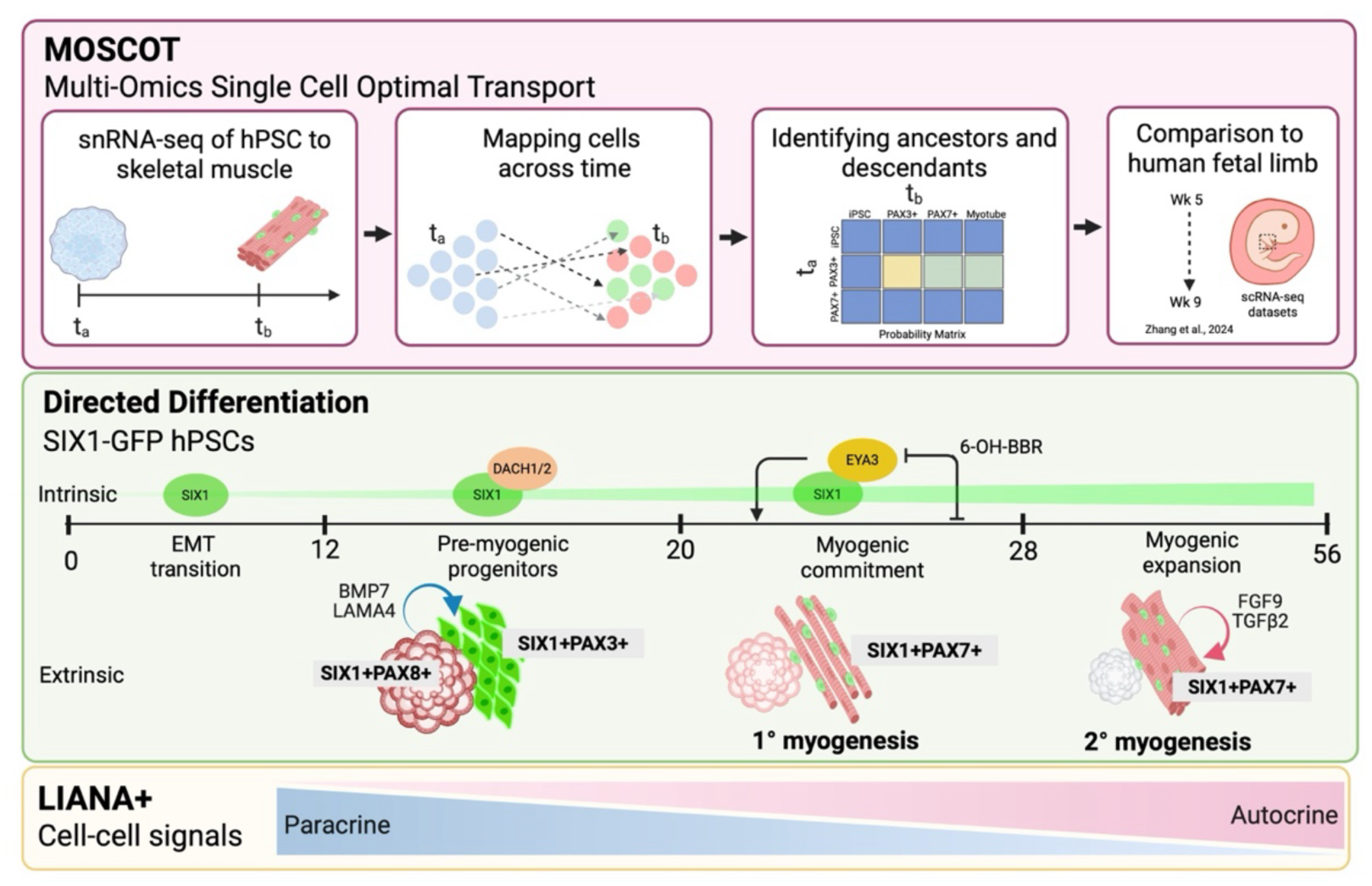

**Highlights:**

- SIX1+PAX8+ niche progenitors promote myogenic differentiation via BMP7- and laminin-dependent signaling.
- Loss of SIX1-EYA3 activity disrupts the pre-myogenic niche and impairs the PAX3-to-PAX7 transition
- Multi-omics single-cell optimal transport resolves previously uncaptured transient intermediates and sequential waves of human myogenesis
- SIX1 lineage tracing identifies CREB5 as a top regulator of the PAX7+ state in human myogenesis

## Introduction

Reconstructing how stem cells transition into differentiated lineages during development and from human pluripotent stem cells (hPSCs) remains a fundamental challenge. Recent advances in single cell transcriptomics have provided valuable insights into the heterogeneity of stem cells during development, enabling a better understanding of disease origin, organogenesis, and lineage specification genes [1, 2]. For skeletal muscle, these approaches these approaches have shown that hPSC-derived skeletal muscle progenitor cells (SMPCs) generated across multiple protocols align with embryonic-to-fetal myogenic stages *in vivo* [3]. Transcriptomic profiling of hPSC-derived SMPCs has also identified cell-surface markers that distinguish progenitor subpopulations with enhanced regenerative potential and molecular signatures relevant to improving cell purification [4–7]. More recently, single-nucleus RNA-seq studies of iPSC-derived skeletal muscle and organoid systems have advanced our understanding of myofiber heterogeneity and regulators of myofiber development but have primarily emphasized endpoint population diversity rather than the transitional biology of lineage commitment [8, 9]. However, human *in vivo* and *in vitro* myogenesis remains incompletely defined as a continuum of growth, and the transient intermediate cell states that precede terminal myotube differentiation. The intermediate states and signaling environments that support human myogenic lineage progression are still poorly resolved.

Optimal Transport (OT), a geometric approach for comparing probability distributions [10], has provided a framework for studying developmental trajectories and inferring ancestor-descendant relationships, with transcriptional programs experimentally validated during hPSC reprogramming [11]. Recently, multi-omics single-cell optimal transport (moscot) has emerged as a powerful approach, incorporating multimodal information to efficiently reconstruct the temporal trajectories of hormone-secreting pancreatic islet cell types during mouse development, with predicted driver genes experimentally validated in hPSC-derived islets [12]. In combination, CellRank [13, 14] adds directionality through RNA velocity, strengthening trajectory inference by predicting terminal states and fate probabilities to reconstruct cellular dynamics. More broadly, integrating OT-based spatiotemporal modalities across large multimodal datasets can better define lineage trajectories and identify transition regulators that may improve lineage-specific hPSC differentiation.

We have previously shown that SIX1 is expressed at all stages of myogenesis, including early PAX3+ progenitors and later PAX7+ progenitors and myotubes [15], making it an ideal factor for studying skeletal muscle development over time and for validating moscot-predicted transitions. Six1 was first characterized as an essential factor for compound eye formation and is now recognized as critical regulator of organogenesis in multiple tissues [16, 17]. Six1 knockout mice exhibit multisystem developmental defects, including reduced or absent kidneys, severe muscle defects, rib cage deformations, and loss of inner ear structures [18]. In the myogenic lineage, Six1 loss impairs Pax3 expression and prevents delamination of muscle progenitors into the limb, leaving limb buds devoid of muscle precursors [19]. Six1 functions within the Pax-Six-Eya-Dach regulatory network, particularly with Eya proteins, whose phosphatase activity provides an important switch for transcriptional control [18, 20]. Co-expression of SIX and EYA proteins activates the myogenin promoter and Myod enhancers, reinforcing Six1’s role as a master regulator of the myogenic regulatory network [21–25]. Although these factors are also expressed during human skeletal myogenesis [26], how SIX1 regulates the emergence of PAX3+ SMPCs, the transition to PAX7+ SMPCs, and subsequent differentiation to myotubes is unclear.

Directed differentiation depends on orderly progression through sequential developmental states, and errors in any intermediate step can cause profound defects or complete failure of hPSC-derived myogenesis. While our system induces mesoderm lineages, much of the differentiation is guided by cell intrinsic and extrinsic cues yet to be defined *in vitro*. In development, trunk and limb muscles depend on proper segmentation of the paraxial mesoderm into embryonic somites which develop in close association with other transient embryonic tissues, including lateral plate and intermediate mesoderm [27]. However, in directed differentiation the extent to which these co-arising transient lineages contribute to skeletal muscle formation is unclear. During embryonic and fetal myogenesis, the dorsal somite, or dermomyotome, gives rise to the earliest SMPCs, marked by Pax3 [28, 29]. Pax3+ progenitors generate early myofibers and transition to Pax7+ precursors, some of which later establish the adult satellite cell (SC) pool required for lifelong regeneration [30, 31]. Skeletal muscle development also depends on hierarchical activation of the myogenic regulatory factors Myf5, MyoD, myogenin, and MRF4; in their absence, muscle fails to form and cells adopt non-myogenic fates [32, 33]. Our hPSC myogenesis system provides a platform to interrogate these programs over time, including how paraxial mesoderm and other mesoderm lineages co-arise *in vitro* to influence myogenic fate progression.

Leveraging an *in vitro* system that enables derivation of multiple mesoderm cell types with high myogenic yields, we set out to resolve the mechanisms of human myogenic lineage commitment through transient developmental states and terminal stages of *in vitro* myogenesis. We combined longitudinal single-nucleus RNA sequencing, moscot, a new OT-based trajectory inference, and a SIX1-GFP reporter for functional lineage tracking of successive myogenic transitions and defining the intrinsic and extrinsic signals governing myogenic progression. Moscot identified SIX1 as a top driver of each myogenic state *in vitro* and *in vivo*, while distinct SIX1 cofactor expression distinguished successive states. Directed differentiation to muscle generated spatially organized 3D structures consistent with a transient SIX1+PAX8+ intermediate mesoderm population temporally linked to SIX1+PAX3+ myogenic progenitors that provided a pre-myogenic niche supporting the PAX3-to-PAX7 transition. Using a SIX1-GFP reporter hPSC line, we further found that PAX3+ cells failed to proliferate in the absence of the PAX8+ niche. LIANA+ analysis implicated a temporally restricted BMP7- and laminin-dependent signaling between PAX8+ niche cells and skeletal muscle progenitors and provided an extrinsic mechanism to promote early myogenesis. Pharmacologic disruption of the SIX1-EYA3 during the key myogenic transition window; however, impaired the PAX3-to-PAX7 transition, reduced myogenic marker expression, collapsed the 3D intermediate mesoderm niche, and enriched for a Desmin+ non-myogenic mesenchymal/fibrotic-like cell. Together, these findings provide a foundation for improving skeletal muscle progenitor production and establish a human *in vitro* platform to define how SIX1 and co-arising niche populations and intrinsic regulatory networks coordinate early myogenic lineage commitment.

## Results

### Optimized single-nucleus profiling and integration to enable moscot-based analysis of hPSC myogenesis

Because hPSC-directed skeletal muscle cultures contain multinucleated myofibers and complex 3D structures, we established a single-nucleus workflow that preserved nuclear integrity across myogenic differentiation and improved sequencing quality metrics to profile all cell types across time. To optimize nuclei isolation, we compared pretreatment with the ROCK inhibitor Y-27632 and the small-molecule cocktail CEPT, which has been shown to improve survival and reduce cell stress [34, 35]. CEPT pretreatment better preserved cellular structure during dissociation, protected nuclei from physical deformation, and enabled fluorescence-activated cell sorting (FACS) to remove debris and doublets before 10x Chromium single-nucleus paired-end RNA-sequencing (**Fig. S1A**). Following this optimization, sequencing quality improved substantially across timepoints, with reduced variability and an acceptable fraction of reads in cells (73–85%; **Fig. S2B**).

We then profiled 124,574 nuclei spanning seven stages of hPSC skeletal muscle differentiation. We captured cells at the start of directed differentiation of hPSCs (day 0), during mesoderm induction (day 5), during SMPC emergence (day 12) and expansion (days 20 and 28), and through their differentiation to myotubes (days 42 and 56) (**Fig 1A**). To integrate these temporally diverse cell states, we compared six computational approaches and selected scVI/scANVI, which best preserved the biologically expected separation of pluripotent and differentiated populations while maintaining coherent progression across time (**Fig S3** and **Extended Methods**)[36, 37]. This optimized workflow provided the foundation for downstream trajectory analysis of hPSC myogenesis that longitudinally tracked 25 transient or stable myogenic and non-myogenic populations across time (**Fig 1B** and GSE323376).

**Figure 1.**
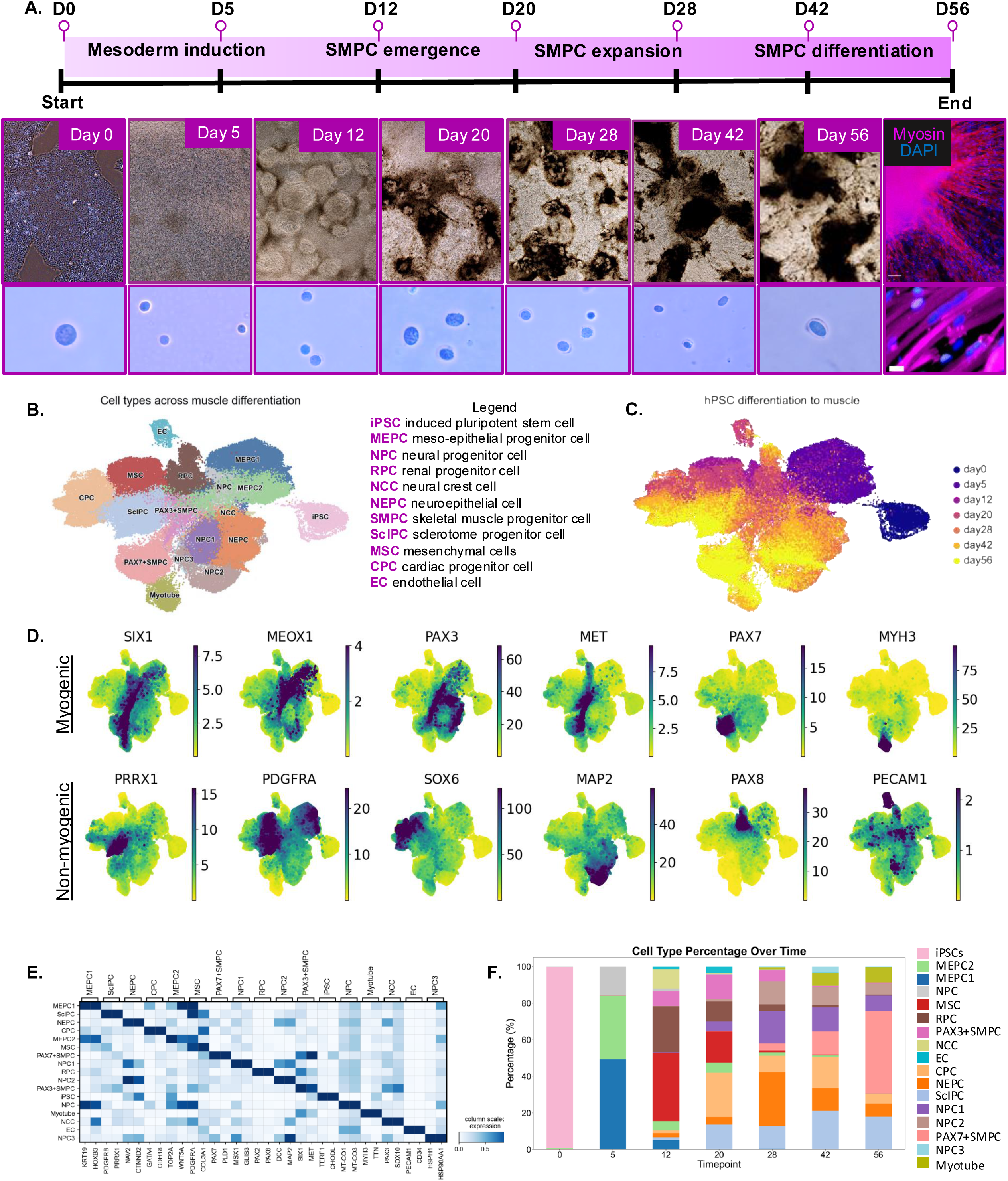
SnRNA-seq identified major cell types emerging from directed differentiation of hPSCs to skeletal muscle. **A.** Timeline of key timepoints during skeletal muscle differentiation protocol taken for snRNA-seq analysis. *Top*: Brightfield images show representative morphologies across time. Scale bar equals 200μm. *Bottom*: Insets of isolated nuclei for snRNA-seq. Scale bar equals 20μm. IF image shows DAPI in blue and MF20 in magenta representing robust myotube differentiation. **B-C.** UMAP plot generated through scVI integration algorithm depicting the cells that arise throughout differentiation protocol, labeled by cell type identity and time. **D.** scVI normalized feature plots showing selected gene marker expression unique to myogenic and non-myogenic cell types. **E.** Matrix plot shows the top 2 DEGS across cell types; values show the mean expression per gene grouped by cell type identity; 1 represents maximum mean expression and 0 is the minimum. **F.** Bar graph depicting cell type proportions (%) over time; colors match cell type annotations on UMAP (also see **TableS1**).

### SnRNA-seq time-course analysis reveals dynamic population changes during hPSC myogenic differentiation

Following mesoderm induction from hPSCs, we identified the emergence of a transient mesodermal progenitor cell with epithelial-like properties (MEPCs) at day 5. MEPCs expressed mesoderm (*MSGN1*) and epithelial (*KRT19*) genes, weakly expressed *MESP2* and *TCF15*, and did not express *UNCX*, suggesting these cells occupy a transition state prior to somitogenesis [38]. However, MEPCs also expressed *HOX* and *MEIS* genes involved in body patterning and branchial arch outside of the somite, indicating a mixed mesodermal identity [39] (**Fig. S4A**).

At later timepoints, we captured diverse myogenic and non-myogenic populations, several of which were marked by *SIX1* expression. These included early *MEOX1*+, *PAX3*+ and *MET*+ SMPCs, and later *PAX7*+ SMPCs, while *MYH3* distinguished differentiated myotubes (**Fig. 1D**). Non-myogenic populations included persistent populations of *PRRX1*+ sclerotome-like progenitor cells (SclPCs), which localized near *PDGFRA*+ mesenchymal-like cells (MSCs) in UMAP space, as well as a distinct population expressing early vascular or cardiac-like progenitor (CPC) genes, including *SOX6* and *GATA4*. Finally, we captured a distinct *PAX8*+ intermediate mesoderm progenitor that progressed toward a renal-like progenitor (RPC) identity (**Fig. 1D**). These lineage-defining markers are summarized in the heatmap and ranked among the top differentially expressed genes for their respective cell types (**Fig. 1E**).

Because integration of multiple time-specific datasets can cause global transcriptional differences to dominate, potentially masking subtle but biologically meaningful variation within individual timepoints [40], we also performed independent timepoint-specific analyses, with particular emphasis on the myogenic lineage (see Extended Methods and **Figs. S4-5**). These analyses confirmed identify of an additional small subset of cells (∼1.4%) at day 5 expressing metabolic and neuronal-associated genes like *MAP2* and *SOX2*, consistent with later populations corresponding to *MAP2*+ neural progenitor cells (NPCs) and neural epithelial cells (NEPCs) (**Fig. 1D-F** and **S4A**).

We found cellular identities shifted rapidly and extensively over time in both integrated datasets and individual timepoint analyses (**Fig. 1F**). Following mesoderm induction, the differentiation landscape expanded quickly from three major populations at day 5 to at least eight distinct populations by day 12, highlighting the striking cellular complexity that emerges early during hPSC myogenesis. In the integrated dataset, MSCs (37%, red) and RPCs (25%, brown) together comprised most cells at day 12, but both populations declined sharply and were only minimally detected by day 28, representing ∼1% and 3% of cells, respectively. In contrast, other populations persisted with relatively modest fluctuation, including SclPCs (light blue), which accounted for 12%–20% of cells from day 20 onward. The myogenic compartment also underwent marked temporal restructuring. PAX3+ SMPCs expanded rapidly from days 12 and 20 (8.0% and 13.3%), decreased by day 28 (6%), and were minimally detected by day 42 (0.5%). Notably, PAX3 and PAX7 displayed a strong inverse relationship over the course of differentiation. As PAX3+ SMPCs expanded between days 12 and 20, PAX7+ SMPCs constituted only ∼0.3% of the population. However, as PAX3+ SMPCs declined, PAX7+ SMPCs increase exponentially, reaching 3.8% by day 28 (peach) and subsequently expanding to nearly half of the total cell population (45%) by day 56 of differentiation. Uniquely enabled by snRNA-seq, we also captured terminally differentiated myotubes beginning at day 28 that reached ∼10% of the population by the end of directed differentiation (**Figs. 1F, S5, and Table S1**).

To assess reproducibility at the end of directed differentiation, we performed snRNA-seq on two independent differentiations and integrated the datasets using scVI. Both replicates yielded similar cell type compositions. At this terminal stage, we identified non-proliferative (23-32%) and proliferative (6.6-7.0%) SMPC subpopulations, distinguished by the absence or presence of proliferation genes such as MKI67 and hyaluronan mediated motility receptor (HMMR) at both the RNA and protein levels. To determine whether cells continued to transition beyond day 50, we combined RNA velocity (scVelo) with CellRank [41] [13]. These analyses showed that most cells had reached terminal-directed trajectories with limited onward transitions, consistent with CellRank’s prediction of 10 terminal macrostates, including non-myogenic populations that co-arise with skeletal muscle at the end of differentiation. Random-walk simulations further indicated that both proliferative and non-proliferative SMPC subpopulations remain myogenic, and predominantly converge on the myotube state (**Fig. S6**).

### Moscot reconstructs the origins of myogenic progeny and successive waves of myotube formation

To define the temporal transitions that underlie human myogenesis *in vitro*, we applied the optimal transport framework with moscot, mapping cells across timepoints to inferring lineage relationships and shared transcriptional states [12]. To assess how well moscot reconstructed myogenic trajectories and to determine how hPSCs progress toward skeletal muscle, we subsetted the analysis on the myogenic lineage by including hPSCs (day 0), all progenitor populations (day 5), PAX3+ SMPCs (days 12-20), and PAX7+ SMPCs and myotubes (days 28-56) (**Fig. 2A**). We then reconstructed lineage progression through sequential timepoint pairs across directed differentiation generating transport matrices that infer ancestor-descendant relationships over time.

**Figure 2.**
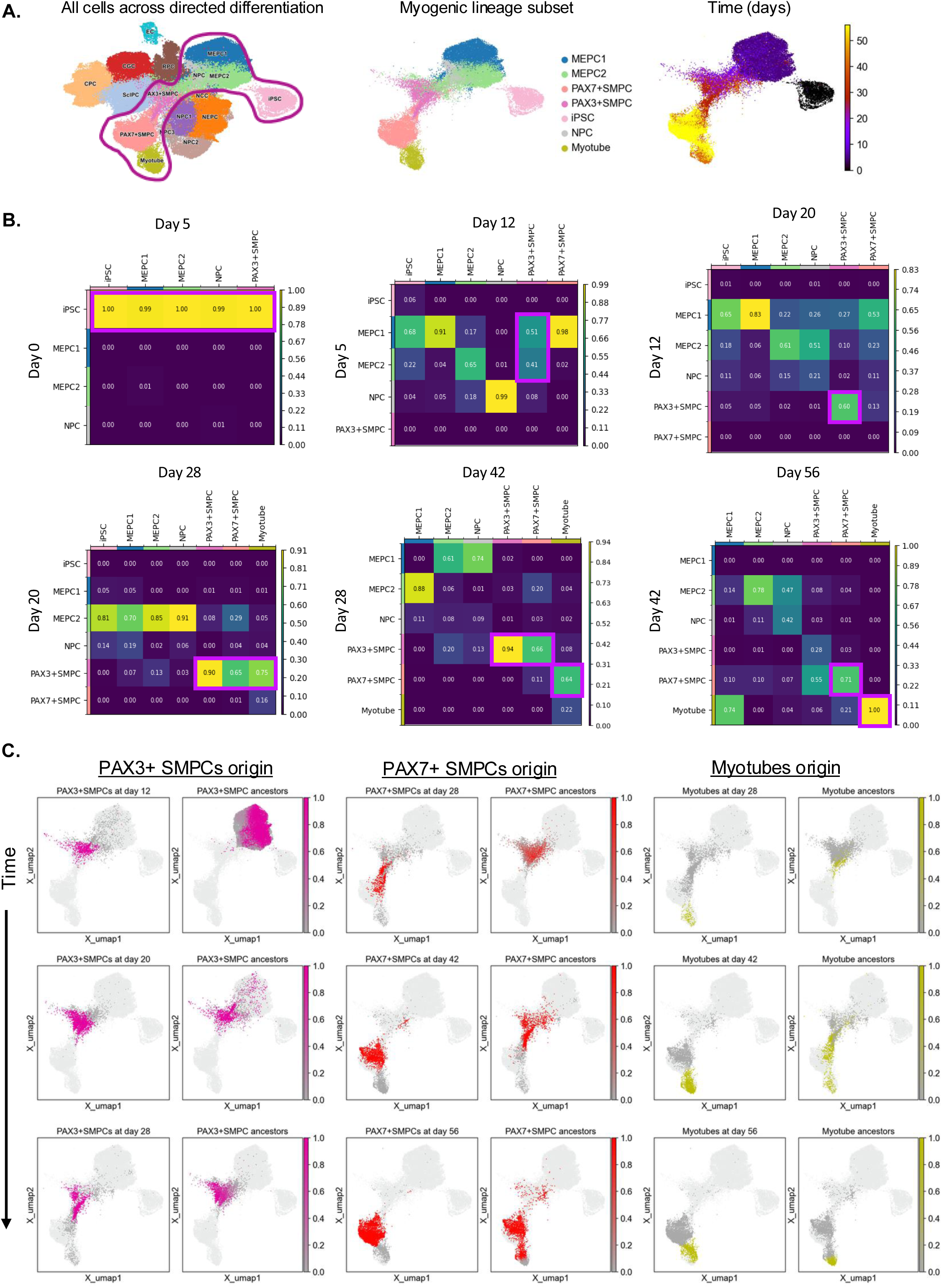
Moscot identifies waves of myogenic progeny giving rise to distinct myotube subsets. **A.** Sub-setting of the myogenic lineage for multi-omics single cell optimal transport (moscot). D0 iPSCs, D5 subtypes, D12-56 PAX3 and PAX7 SMPCs, and myotubes extracted & labeled by cell type identity and time. **B.** Reconstruction of myogenic trajectories highlighted by the transition matrices showing the ancestry of each cell type, modeled by the temporal problem in the moscot algorithm. iPSC induction days 0-5; PAX3 derivation from MEPCs days 5-12; PAX3 expansion days 12-20; primary myogenesis by PAX3+SMPCs days 20-28; secondary myogenesis by PAX7+SMPCs days 28-42, myogenic expansion days 42-56. **C.** UMAP visualization of PAX3+SMPCs, PAX7+SMPCs, and myotube descendants across time. The right plot shows the ancestry of each cell at a specific time.

Moscot predicted that all day 5 populations (MEPC1, MEPC2, and NPCs) arose from hPSCs, as expected (prob = 1). From day 5 to day 12, MEPC1 (prob = 0.51) and MEPC2 (prob = 0.41), but not NPCs (prob = 0.08), contributed to the emergence of SIX1+PAX3+ SMPCs. At day 12, PAX3+ SMPCs comprised ∼8% of the population, whereas PAX7+ SMPCs were nearly absent (0.01%) (**Fig. 1F**), suggesting that any predicted early contribution of MEPC1 to PAX7+ SMPCs is likely overstated by the model. This interpretation was supported by the day 12 to day 20 transition matrix, in which PAX3+ SMPCs primarily gave rise to themselves (prob = 0.60). From day 20 to day 28, however, PAX3+ SMPCs strongly contributed to the emergence of PAX7+ SMPCs (prob = 0.65), consistent with the onset of the PAX3-to-PAX7 transition.

Interestingly, moscot also resolved two distinct waves of myotube formation. Between days 20 and 28, PAX3+ SMPCs were the dominant source of MYH3+ myotubes (prob = 0.75), marking an initial wave of myogenesis. Between days 28 and 42, myotubes were instead derived predominantly from PAX7+ SMPCs (prob = 0.64), consistent with a second wave of myogenesis. By the end of directed differentiation, PAX3+ SMPCs, PAX7+ SMPCs, and myotubes largely exhibited autologous self-contribution, indicating stabilization of their terminal identities (**Fig. 2B**).

The lineage relationships were also visualized on UMAP projections (**Fig 2C**). UMAPs highlight the emergence of PAX3+ SMPCs at day 12 from corresponding day 5 MEPC ancestors (pink). Over time, PAX3+ SMPCs initially sustained their own population and then gave rise to PAX7+ SMPCs between days 28 and 42 (red). Similarly, the earliest myotubes detected at day 28 were predicted to arise from PAX3+ cells, whereas myotubes at day 42 were primarily linked to PAX7+ SMPCs (green). These data suggested that hPSC-directed myogenesis recapitulates successive developmental waves *in vitro* consistent with embryonic and fetal myogenesis *in vivo*.

### Transcription factor inference identifies candidate regulators of temporal myogenic state transitions

Using the predicted origin distributions revealed by moscot, we also identified the top transcription factors involved in ancestor/descendant transition matrices for early, intermediate, and late drivers of myogenesis. Using the lineage relationships extracted with moscot, we predicted myogenic cell fate probabilities and revealed transition-specific driver genes involved at each transitional step (**Fig. 3**). We then cross-referenced the expression of our unique transcription factor inferences using a human embryonic limb cell atlas [2].

**Figure 3:**
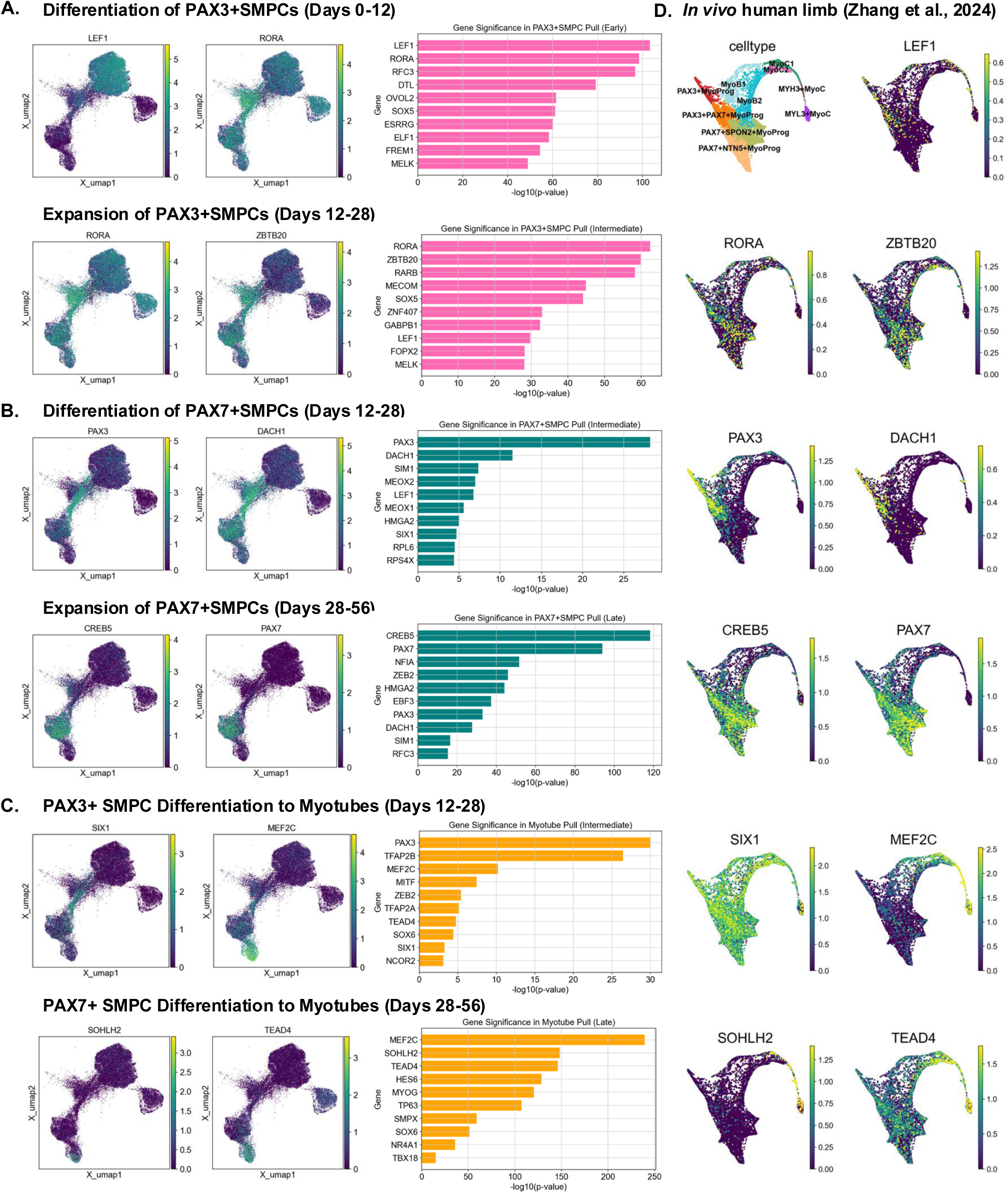
Transcription-factor inference highlighted putative drivers of the PAX7+ states overtime. **A.** UMAP depicting expression of moscot’s top correlated transcription factors computed using the compute correlation of push forward or pull-back distribution (see moscot documentation) of PAX3 SMPCs between days 0-12 or 12-28, **B.** PAX7 SMPCs between days 12-28 or 28-56, and **C.** myotube differentiation between days 12-28 and 28-56. Bar graphs on the right depict log transformed p-values correspond to the top 10 transcription factors following moscot analysis (also see **TableS2**). **D.** Corresponding feature plots of *in vitro* transcription factor marker expression in human limb development.

Among the top candidate regulators of early PAX3+ SMPC differentiation were LEF1, a WNT pathway component implicated in somite myogenesis [42], and *RORA*, which has been linked to metabolic regulation during mesodermal differentiation [43]. Consistent with moscot predictions, *LEF1* was strongly expressed in human *PAX3*+ myogenic limb progenitors *in vivo* (**Fig. 3A**). In agreement with the transition matrices, the top factors associated with early PAX7+ SMPC specification included *PAX3* and *DACH1* (p = 3.22E-12), a *SIX1* cofactor and transcriptional repressor, both of which were also strongly expressed in PAX3+ limb progenitors *in vivo*. Moscot also identified *CREB5* (p = 4.8E-119) as a top regulator of the PAX7 state from days 28–56. Although previously linked to pharyngeal myogenesis [44], our analysis implicates *CREB5* as the number one gene regulating the *PAX7* transition during human myogenesis. Along with *PAX7*, *CREB5* was strongly expressed across PAX7+ precursors *in vivo*, supporting a potentially conserved role in embryonic limb myogenesis (**Fig. 3B**). Strikingly, the earliest PAX3+ SMPC-derived myotubes were associated with distinct candidate regulators, including *TFAP2A, TFAP2B*, and *MITF* (p < 3.6E-08). In contrast, later PAX7+ SMPC-derived myotubes were associated *TEAD4* and *MYOG* (p<3.3E-121), which were selectively expressed in limb myocyte populations *in vivo* (**Fig. 3C**). *MEF2C* was also identified as a top transcriptional driver for both PAX3- and PAX7-associated myogenic differentiation (p<1E-10). Together, these analyses identified candidate regulators of PAX3+ and PAX7+ SMPC specification, factors associated with maintenance of established myogenic states, and distinct drivers of myotube differentiation arising from PAX3+ versus PAX7+ progenitors.

We also found that *SIX1* ranked among the top transcription factors associated with specification of PAX7+ cells from hPSCs and robustly marked myogenic intermediates across differentiation, including PAX3+ progenitors, PAX7+ progenitors, and MYH3+ myotubes. *PAX3* and *SIX1* were found to be in the top ten specification factors for both *PAX7* and myotube transition states (**Fig. 3**). Visualization of expression strength and directionality across the full dataset showed that *SIX1* remained highly specific to SMPCs and myotubes, whereas *PAX3* was lost during myotube differentiation and was also expressed in neuronal subtypes (**Fig. S7A**). Based on the persistent and lineage-restricted expression of *SIX1* across myogenic time, we next used its expression pattern to functionally investigate how SMPC transitions are temporally and spatially regulated during hPSC myogenesis.

### SIX1 lineage tracking captures a developmental window of niche-dependent myogenic progression

To functionally track successive waves of hPSC myogenesis, we used a SIX1:H2B-GFP reporter line [45] across the developmental window spanning the PAX3-to-PAX7 transition (days 20–28) and the subsequent phase of myogenic progression (days 28-42) (**Fig. 4A**). GFP expression emerged by approximately day 7 and persisted throughout directed differentiation, enabling prospective isolation of SIX1+ cells at multiple stages (**Fig. 4B**). Across five independent differentiations, ∼20% of cells were GFP+ at days 20 and 28, increasing to ∼40% by day 42; an intermediate GFP population was observed specifically at day 20, consistent with a transient developmental state (**Fig 4C**).

**Figure 4.**
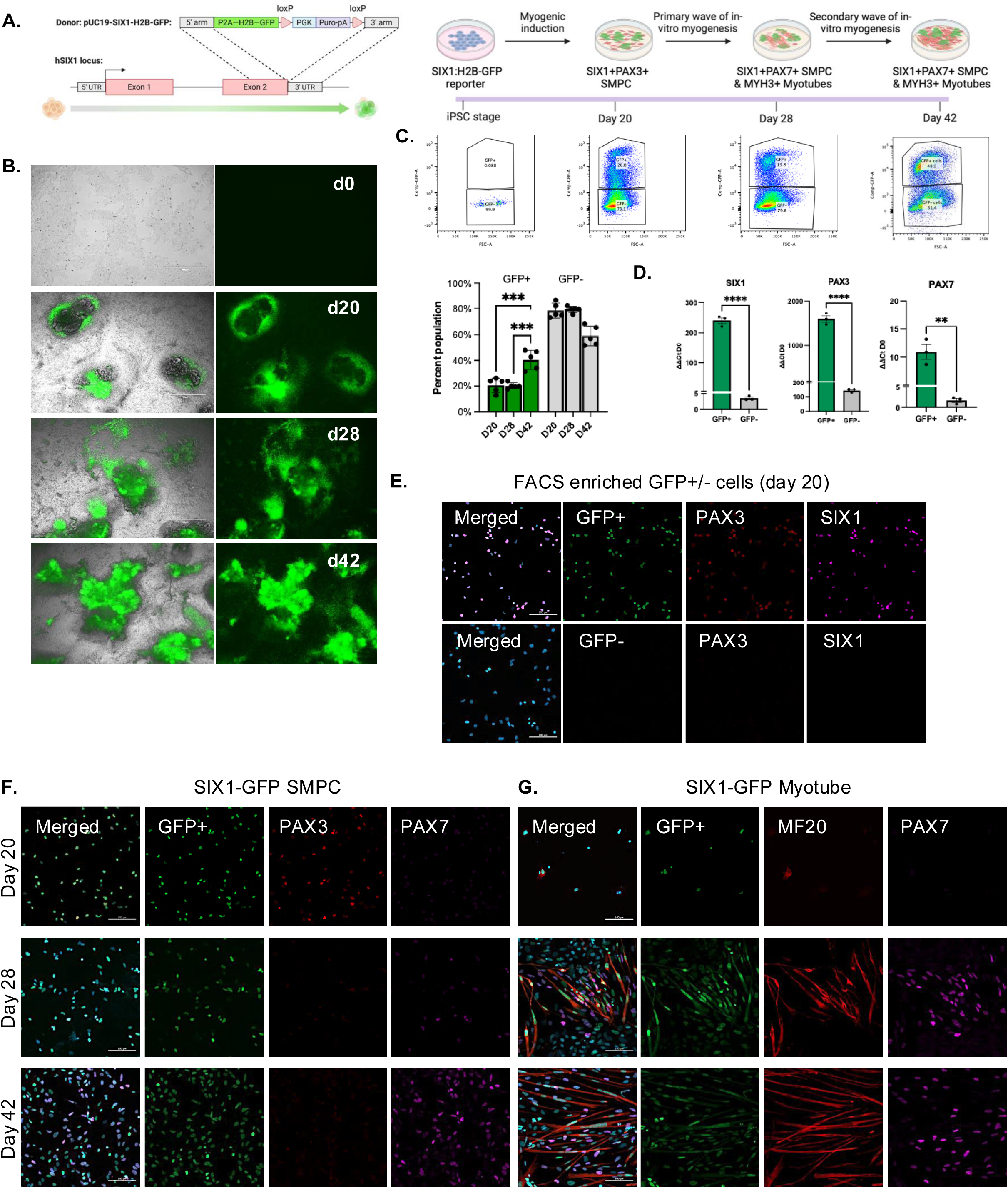
Capturing myogenic waves of in-vitro SMPCs through functional analysis of a SIX1:H2B-GFP reporter hPSC. **A.** Schematics of SIX1:H2B-GFP reporter line and experimental design. **B.** Brightfield and fluorescent images of SIX1-H2B-GFP hPSCs at days 0-42 of directed differentiation. Scale bar equals 750μm **C.** Flow cytometry plots depicting percent expression of H9 and SIX1-GFP+ SMPCs on days 20-42 and corresponding quantifications. (n=15 independent wells, 3 wells combined per sample (n=5)). One-way ANOVA statistical analysis (***P<0.001). **D.** Quantitative rt-qPCR expression of *SIX1, PAX3*, and *PAX7*. Data normalized to GAPDH and relative to expression at day 0 (n=3 biological replicates). t-test statistical analysis (***P<0.001, **P<0.01). **E.** Immunofluorescent images of day 20 GFP+/− FACs-sorted cells stained with DAPI (blue), endogenous GFP (green), PAX3 (red), and SIX1 (magenta). **F-G.** Immunofluorescent images of days 20-42 SIX1-GFP+ SMPCs and myotubes stained with DAPI (blue), endogenous GFP expression (green), PAX3 (red), and PAX7 (magenta). Scale bar equals 100μm.

Immediately after sorting, GFP+ and GFP- fractions were assayed for myogenic identity. At day 20, the GFP+ fraction showed >200-fold enrichment of *SIX1*, with significant enrichment of both *PAX3* and *PAX7*, although *PAX3* expression predominated (p<0.05) (**Fig. 4D**). Endogenous GFP signal overlapped with SIX1 protein expression and was absent from the GFP- population (**Fig. 4E**). Protein expression further validated the presence of PAX3 expression at day 20 but not at later stages, whereas PAX7 was absent at day 20, weakly detectable at day 28, and strongly expressed by day 42 (**Fig. 4F**). Neither PAX3 nor PAX7 were detected in the GFP- fraction (**Fig. S7B**).

To test developmental competence, sorted SIX1+GFP+ cells were replated under myogenic differentiation conditions. Strikingly, day 20 SIX1+ cells failed to survive, proliferate, and differentiate when isolated from the heterogenous niche, indicating that they had not yet acquired autonomous myogenic potential. In contrast, day 28 SIX1+ cells expanded and formed primitive myotubes, and by day 42 they generated robust myotubes (**Fig. 4G**). These findings indicate that day 20 myogenic progenitors remain dependent on a co-arising pre-myogenic niche for survival and expansion, whereas between days 20 and 28 they undergo a key developmental transition that enables committed myogenic progression.

### SMPCs rely on extrinsic paracrine and autocrine signals to achieve myogenic commitment

The failure of day 20 SIX1+PAX3+ progenitors to survive and proliferate after FACS isolation suggested that these cells remain dependent on a transient co-arising niche population, prompting us to investigate the cell types that were most abundant during the early stages of differentiation. Notably, our snRNA-seq analysis showed that SIX1 gives rise to two divergent populations at day 12: a SIX1+PAX3+ paraxial mesoderm-derived SMPC and SIX1+PAX8+ intermediate mesoderm-derived RPC (**Fig. 5A**). The PAX8+ population was most prominent at early timepoints, consistent with a transient niche population present during the window of myogenic commitment. This is also consistent with *in vivo* studies showing Six1 functions in both paraxial mesoderm-derived skeletal muscle development [46] and intermediate mesoderm-derived kidney development intermediate mesoderm derived kidney development [47]. Immunofluorescence revealed that SIX1+PAX8+ cells (pink box) formed tube-like three-dimensional structures, adjacent to where SIX1+PAX3+ SMPCs frequently emerged (green box), also consistent with the proximity of intermediate mesoderm that develops between the somites and the lateral plate mesoderm *in vivo*. SIX1+PAX3+ SMPCs remained closely associated with these 3D structures but observed that SIX1/PAX3 expression shifted over time, from the core of the structure to neighboring cells (**Fig. 5B**), revealing a dynamic spatial pattern of SIX1 activation during early myogenic progression.

**Figure 5.**
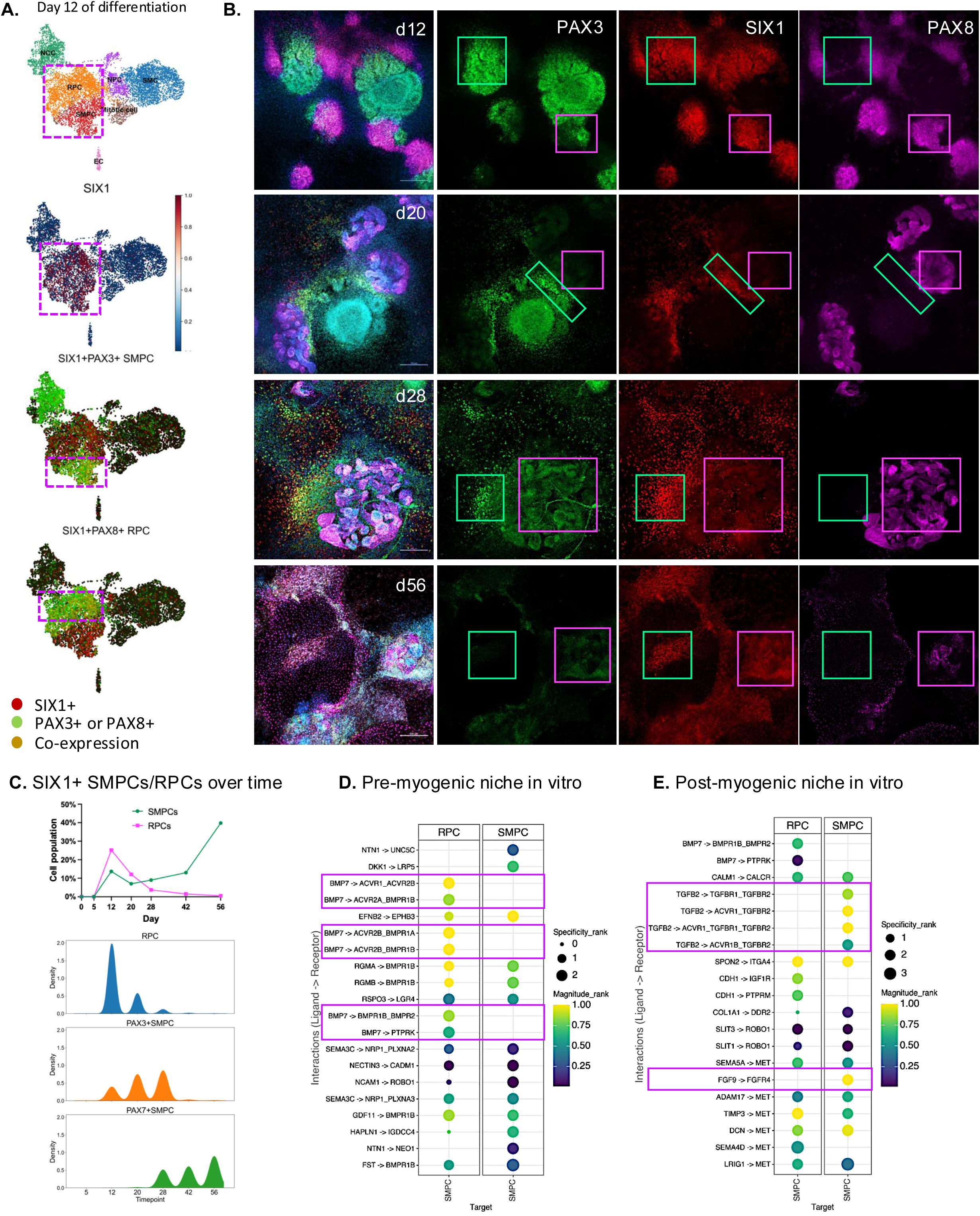
Intrinsic and extrinsic signals of PAX7+ SMPC lineage commitment. **A.** Feature plot depicting expression of SIX1 by PAX3+ SMPCs and PAX8+ RPCs at day 12. **B.** Immunofluorescent images of days 12, 20, 28, and 56 directed differentiation cultures stained with DAPI (blue), PAX3 (green), SIX1 (red), and PAX8 (magenta). Green box highlighting PAX3+SIX1+ cells and magenta highlighting PAX8+SIX1+ cells. Scale bars: 200um. **C.** Line graph depicting snRNA-seq percent population (percent total) of SMPCs and RPCs across time in differentiation. Density plot reflecting gene expression of SIX1 by RPCs, PAX3+SMPCs, and PAX7+SMPCs across time. **D-E.** Dot plot showing LIANA’s rank aggregate prediction of top 20 ligand-receptor interactions between RPCs and SMPCs ranked by magnitude in the pre-myogenic niche (day 12) and post-myogenic niche (day 28).

To visualize SIX1 expression patterns in RPCs and SMPCs across time, we generated density plots from our integrated dataset. At day 12, RPCs exhibited the highest levels of SIX1 expression relative to SMPCs; but this expression declined sharply over time. In contrast, PAX3+ SMPCs maintained relatively stable expression, while *SIX1* emerged in PAX7+ SMPCs by day 28 through the end of the derivation (**Fig. 5C**). Together, these data suggest that dynamic SIX1 expression marks lineage divergence between PAX8+ and PAX3+ derivatives and support a model in which early SMPCs rely on a transient PAX8+ intermediate mesoderm niche for survival and progression through myogenic commitment.

To define cell-cell interactions within this pre-myogenic niche, we used LIANA+ to infer steady-state ligand-receptor communication [48] between RPCs and SMPCs before commitment (days 12–20) and after commitment (day 28). Ranking the top interactions by consensus strength revealed that prior to myogenic commitment, SMPCs interacted strongly and specifically with RPCs through the *BMP7-BMPR1B* ligand-receptor pair (**Fig. 5D**). Although *BMPR1B* was not exclusive to SMPCs, BMP7 expression was restricted to RPCs in UMAP space. Multiple *BMP* and *SEMA3* ligand subtypes were also among the top interactions between RPCs and SMPCs (**Fig. S8A**). In addition, LIANA+ identified RPC-derived laminins (*LAMA1, LAMC1*, and *LAMA4*) with strong *DAG1* interactions on SMPCs, indicating that the pre-myogenic niche provides both secreted and extracellular matrix support (**Fig. S8B**).

As SMPCs commit to the PAX7+ myogenic lineage at day 28, LIANA+ identified several autocrine interactions specific to the myogenic compartment (**Fig. 5E** and **S8C**). Among these ligand-receptor pairs were *FGF9-FGFR4, TNC-ITGA5*, and *TGFB2-TGFBR2* that were exclusively expressed by myogenic cells. SMPCs also continued to receive autocrine and paracrine input through *MET* and *ROBO1*, whereas RPC-derived BMPs, laminins, and other growth factors became progressively reduced and were no longer detected at later stages as the RPC population declined. Thus, these data indicate that SMPCs are closely linked to a transient SIX1+PAX8+ intermediate mesoderm population and that this co-arising niche provides extrinsic signals required for survival and progression through early myogenic commitment.

### SIX1+ progenitors transition from a niche-dependent pre-myogenic state to committed myogenic cells

To define genes associated with acquisition of myogenic competence, we compared SMPC transcriptomes with all other cell types at their respective timepoints. We found Day 20 and day 28 SMPCs shared 418 differentially expressed genes (DEGs; adjusted p<0.05), including canonical developmental regulators such as *PAX3, SIX1/4, EYA1/2/4, MEOX1/2,* and *MET*, consistent with a shared embryonic myogenic program [49] However, each stage also displayed a distinct transcriptional profile, with 521 unique genes at day 20 and 964 at day 28 **(Fig. 6A**).

**Figure 6.**
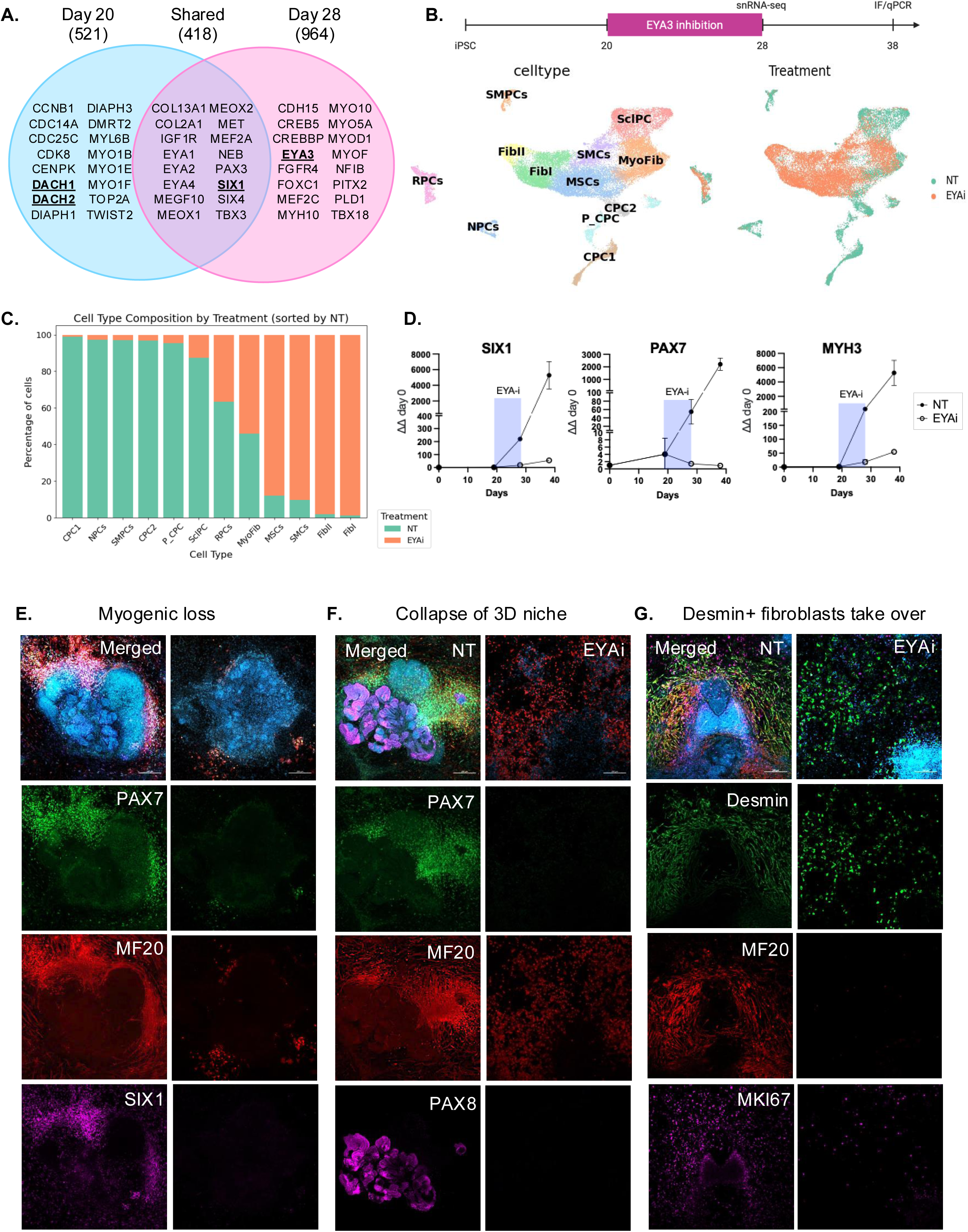
Inhibition of SIX1/EYA3 between days 20-28 prevents myogenic and non-myogenic differentiation. **A.** Venn Diagram showing selected unique and shared DEGs between day 20 and day 28 SMPCs (also see **TableS3**). **B.** Schematic of experimental layout, and UMAP post integration of non-treated and 6-hydroxybenzbromarone-treated (EYAi) differentiations at day 28, labeled by cell type identity and treatment. **C.** Stacked bar graph showing cell type composition by treatment and sorted by highest to lowest percentage of cells in non-treated cells (see **TableS6** for cell type proportions). **D.** Quantitative RT-qPCR shows levels of *SIX1, PAX7*, and *MYH3* in non-treated and EYAi treated cultures relative expression to GAPDH and normalized to hPSCs at day 0. **E-G.** Immunofluorescent images of non-treated (NT) and EYAi-treated directed differentiation cultures at day 28 stained with **E.** DAPI (blue), PAX7 (green), MF20 (red), and SIX1 (magenta). **F.** DAPI (blue), PAX7 (green), MF20 (red), and PAX8 (magenta). **G.** DAPI (blue), PAX3 (Desmin), MF20 (red), and MKI67 (magenta). Scale bar equals 200μm.

At day 20, SMPCs were enriched for genes linked to cell-cycle regulation and expressed several non-muscle myosin family members, consistent with a proliferative but not yet fully committed state. By day 28, SMPCs upregulated genes linked to skeletal muscle identity and differentiation, including *CDH15, FGFR4, MEF2C, MYOD1*, and *CREB5*, the latter identified by moscot as a top regulator of the PAX7 state. Notably, the SIX1 cofactor network also shifted across this window, with *DACH1/DACH2* enriched at day 20, whereas the SIX1 co-activator *EYA3* was upregulated at day 28.

Pathway-level analysis using the Human Molecular Signatures Database in EnrichR [50] further supported this transition. Day 20 SMPCs were most strongly associated with cell-cycle and mitotic programs, whereas day 28 SMPCs were enriched for gene sets linked to differentiation, epithelial-to-mesenchymal transition, and myogenesis (**Fig. S9A** and **Extended Methods**). Together, these data indicate that between days 20 and 28, SIX1+ progenitors transition from a pre-myogenic, niche-dependent state to a more committed PAX7+ myogenic state, and that this shift is accompanied by dynamic changes in SIX1 cofactor interactions, most prominently the transition from DACH1/2 to EYA3.

### Inhibition of SIX1/EYA3 prevents the myogenic transition from PAX3 to PAX7 and disrupts the pre-myogenic niche

To investigate how SIX1 signaling directs myogenic differentiation, we focused on its cooperative interactions with the EYA family of co-factors. Although SIX1-cofactor interactions are known to influence lineage specification in other developmental contexts [51], whether EYA3 functionally contributes to PAX7 myogenic lineage commitment from hPSCs is unknown. To test this, we treated directed differentiation cultures with 6-hydroxybenzbromarone, a EYA3 phosphatase inhibitor (EYAi), during the critical PAX3-to-PAX7 transition window (day 20-28). We then assessed the consequences by snRNA-seq immediately after treatment, and by immunofluorescence and rt-qPCR ten days later (day 38), to determine the broader impact on lineage progression (**Fig. 6A**).

At the end of the treatment, snRNA-seq captured 12 myogenic and non-myogenic populations (**Fig. 6B**). However, EYAi-treated cultures showed a marked expansion of fibrotic-like cells rarely present under normal culture conditions (**Fig. 6C**). These fibroblasts were distinguished by high Desmin expression, a structural protein found in muscle and mesenchymal cells (**Fig. S9B**). In parallel, RT-qPCR revealed an immediate reduction in key myogenic markers, including *SIX1, PAX7*, and *MYH3* (**Fig. 6D**), and protein expression of myogenic lineage markers remained reduced at day 38 relative to untreated controls (**Fig. 6E**).

The effects of EYA3 inhibition were not limited to the myogenic lineage. Morphologically, disruption of SIX1-EYA3 signaling caused a striking collapse of the 3D niche structures, which we now identify as being composed of PAX8+ intermediate mesoderm RPCs (**Fig 6F**). However, these conditions created a permissive environment for Desmin+MF20- cells, as Desmin co-expression with sarcomeric myosin (MF20) was only found in the untreated cultures, suggesting a diversion from terminal myogenic differentiation (**Fig. 6G**). We also observed loss of MAP2+ neuronal progenitors (**Fig. S9C)** and a clear reduction in GATA4+ cardiac progenitors (**Fig. S9D**), indicating broader disruption of lineage specification within the differentiation system. Together, these findings identify SIX1-EYA3 as a central developmental node linking intrinsic myogenic progression to maintenance of the transient PAX8+ pre-myogenic niche and show how disruption of this coupling reshapes the trajectory of human myogenesis.

## Discussion

Our study traces the cellular and regulatory basis of early human myogenesis *in vitro* and identifies a previously unrecognized developmental niche required for myogenic lineage progression. We show, for the first time, that intermediate mesoderm and paraxial mesoderm can be derived in parallel from a human iPSC culture system, and that a SIX1+PAX8+ intermediate mesoderm cells form a transient 3-dimensional pre-myogenic niche that supports SIX1+PAX3+ paraxial mesoderm-derived progenitors to transition into the skeletal muscle lineage through a BMP7- and laminin-dependent signaling. This finding is particularly significant because these transient embryonic tissues are spatially linked during development *in vivo* yet, had not been previously modeled in a human *in vitro* system, limiting investigation of how neighboring lineages coordinate lineage commitment and how their disruption may contribute to multi-organ developmental disorders. Using optimal transport, we longitudinally tracked hPSCs across myogenic directed differentiation and demonstrate that moscot reconstructs hPSC progression through an *in vitro* analogue of transient mesoderm-epithelial cell intermediates to form PAX3+ to PAX7+ skeletal muscle progenitors and myotubes, closely mirroring the myogenic patterns observed in embryonic to fetal *in vivo* development [2, 52]. Functional lineage monitoring with a SIX1-GFP reporter hPSC validated these predicted myogenic waves and revealed a transcriptional shift from uncommitted progenitors to committed myogenic cells. We further defined intrinsic and extrinsic features of this transition, showing that SIX1 cofactors shift from a repressive DACH1/2 state to a permissive EYA3 state. These findings establish a human model for studying how transient mesodermal populations interact to control early myogenic development and provide mechanistic insight into how disruption of this coordination can alter both muscle and non-muscle lineage formation.

Directed differentiation from hPSCs offers a unique platform for studying embryonic cell fate transitions in a human system. Because we direct mesoderm differentiation without early suppression of other mesodermal lineages [53, 54], we recreate part of the transient mesodermal environment *in vitro*. This is developmentally significant because paraxial mesoderm patterning occurs within a shared signaling environment and arises near intermediate mesoderm and other somite derivatives, rather than in isolation [55]. Similarly, in our system SIX1+PAX3+ SMPCs emerge adjacent to duct-like SIX1+PAX8+ 3D structures which secrete pro-myogenic signals like BMP7 and Follistatin, previously shown to expand Pax3-positive progenitors during embryonic muscle growth and delay premature differentiation [56, 57]. BMP signaling has stage-specific roles in skeletal muscle that must be tuned to balance progenitor maintenance, growth, and differentiation during prenatal development [58]. Thus, skeletal muscle commitment is reliant not only on cell-autonomous cues but also on extrinsic signals from neighboring non-myogenic populations that co-arise *in vitro*.

Directed differentiation of hPSCs can reveal mechanistic insights on when and how SIX1-dependent myogenic development may be dysregulated, contributing to disease. SIX1 pleiotropy is clinically evident in Branchio-Oto-Renal (BOR) syndrome, a developmental disorder characterized by hearing loss, skeletal muscle defects, and renal abnormalities and where mutations disrupting the SIX1-EYA1 interaction account for nearly half of the cases [59–61]. Similarly, *in vitro*, we find that SIX1 is not only restricted to the myogenic lineage over time but also marks a PAX8+ renal-like epithelial cell early in differentiation, suggesting that SIX1+ cells are progenitors of multiple and transient lineages within a single platform. Although *SIX1* is needed during early embryogenesis, its activation in adult tissues promotes a neoplastic phenotype [62]. Our studies find that SMPCs adopt a highly proliferative transcriptome prior to committing to the myogenic lineage and highlight a switch from *SIX1* repression by *DACH* to *SIX1* activation by *EYA3*, supporting that modulation of the SIX1-complexes may suppress aberrant cell behavior [63].

Our results pinpoint pivotal timeframes for stage-specific myogenic commitment and differentiation from hPSCs. Functional interrogation with a SIX1+GFP reporter line provided evidence that SMPCs acquire myogenic potential overtime as early SIX1+GFP+ cells fail to proliferate and differentiate, whereas later SMPCs effectively expand and generate myotubes. Through moscot, we identified some of the earliest transcriptional drivers for SMPCs including *PAX3*, *DACH1*, and *SIM1*, whereas after SMPCs entering a myogenic program transcriptional drivers change to *CREB5, PAX7,* and *NFIA*, among others. *CREB5* was our number one significant gene and is an understudied transcription factor in myogenesis. However, our investigations in the human limb atlas dataset found that *CREB5* is highly and uniquely expressed by human fetal week 9 PAX7+ progenitor cells [2] which correlates with the immature PAX7 SMPCs generated by hPSCs [3]. *CREB5* is bZIP transcription factor that is unique from other CREB family proteins as it is not a direct activator of cAMP but has been shown to activate MET in non-myogenic contexts [64]. Our longitudinal capture of myogenic differentiations by snRNA-seq enabled the tracking of PAX3 and PAX7 SMPCs that result in transcriptionally distinct myotubes. The first wave of myotubes were driven by a network of PAX3+ SMPCs through *TFAP2-SIX1-ZEB2* transcription factors whereas a second wave is driven by PAX7+ SMPCs through *MYOG-TEAD4-HES6*. There are many differences in the primary and secondary myofibers that arise in dish, notably, the *TFAP2A/B* genes are gone in later differentiation underscoring heterogeneity and a completely understudied area. Collectively, these results suggest that differential SIX1-cofactor interactions may modulate downstream transcriptional programs that establish a permissive window between myogenic commitment and alternative fates. Disruptions of the SIX1-EYA3 interaction during this central time when SMPCs acquire myogenic commitment led to the collapse of the *in vitro* niche, indicating that SIX1–EYA activity sustains the morphogenetic signals that precede robust myogenesis and offers a framework for dissecting SIX1 regulation of myogenic and non-myogenic lineages.

Rapid multilineage specification before myogenic commitment is a defining feature of early hPSC differentiation in our system. At day 5, we capture a dynamic transcriptomic state that is not yet definitive of somite origin [65], but instead expresses broad embryonic developmental genes like *WNT5A*, *KRT19, MEIS1,* resembling proximal and distal mesenchymal programs in the developing human embryonic limb [66]. In line with this plastic state, the mesenchymal-epithelial progenitor cells (MEPCs) give rise to at least seven distinct lineages early in differentiation. A major feature of this stage is the epithelial-mesenchymal remodeling, including EMT and its reverse process MET, which shape the body plan and dictate differentiation of multiple lineages. During skeletal muscle development, the ventral epithelial somite undergoes EMT to give rise to sclerotome-derived tissues including bones, cartilage, and tendons, whereas the dorsal side retains its epithelial structure and forms the dermomyotome [49, 67]. Consistent with this biology, our analysis identified several regulators of EMT including TGF family components *CDH1*, *SNAI1*, *SNAI2*, ß-catenin, and *ZEB1/2* [68–72]. Our cultures therefore appear to pass through *bona fide* EMT intermediates en route to skeletal muscle while allowing a permissive environment for multiple lineage differentiation. When properly established, this unique development niche supports the long-term maintenance of PAX7+ SMPCs for several months *in vitro*.

In this study, we designed a longitudinal snRNA-seq resource that captures large, multinucleated myotubes with numerous fragile and transient cell populations. Because hPSC-derived cultures present substantial technical challenges for nuclei isolation and integration across highly distinct developmental states, we optimized both sample preparation, including CEPT treatment to reduce ambient RNA contamination, and downstream analysis using scIV integration. Our approach enabled robust reconstruction of human myogenic progression while minimizing technical noise that often can complicate interpretation. To support broader use of this resource, we have also made the dataset publicly available (GSE323376) together with a GitHub repository containing 16 custom analysis scripts that enable users to explore and reuse the data more readily.

Through implementation of moscot trajectory analysis, we have better defined myogenesis *in vitro* and related it to human limb development *in vivo*. These studies identify how hPSCs acquire myogenic commitment and point to modulation of the SIX1-EYA transcriptional complex as a potential strategy for improving myogenic differentiation and understanding developmental disease. An important next question is how SIX1 co-regulation selectively favors the myogenic lineage and why SIX1+ progenitor intermediates are required for successful lineage progression. Given that musculoskeletal disorders are a leading cause of global disability by an aging population [73], the ability to direct and standardize human myogenic trajectories is likely to accelerate preclinical modeling for regenerative medicine.

## Acknowledgements

We thank all members of the Hicks lab for their support. Thank you to Drs. Marius Lange and Dominik Klein who provided advice on moscot analyses. We thank Drs. Jason Tchieu and Lorenz Studer for providing the SIX1:H2B-GFP hPSC lines. Thank you to Drs. Edwin Monuki, Marcus Seldin, and Peter Donovan for their mentorship. A special thanks to Hawra Karim for helping to optimize nuclei isolation and Victoria Espericueta for providing CEPT reagents. Sequencing libraries were prepared by Quy Nguyen, Christina Lin, and Melanie Oaks at the UCI Genomic High Throughput Facility. We thank the UCI stem cell research center flow cytometry core Vanessa Scarfone and Pauline Nguyen for assistance with cell sorting. M.R.H. is funded by NIH/NIAMS R01AR084027, O.G.J. is funded through a CIRM predoctoral fellowship EDUC4-12822 and UC President’s Dissertation Fellowship.

---

**Supplemental Figure 1.**
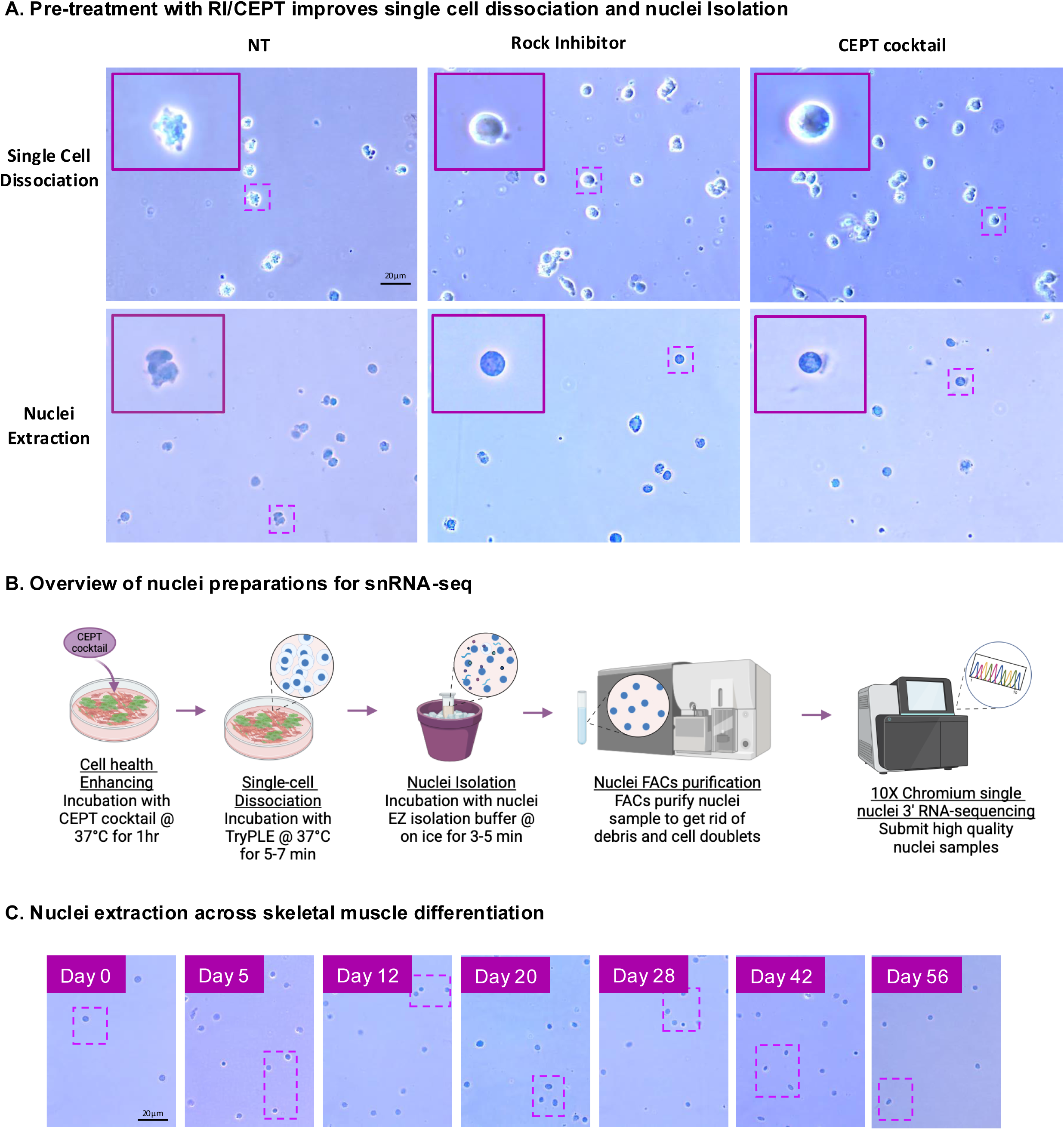
Supplementation with CEPT cocktail prior to single cell dissociation dramatically enhances nuclei integrity. **A.** Representative brightfield images (40X EVOS XL) of cells (top) and nuclei (bottom) treated with and without ROCK inhibitor and CEPT cocktail prior to single cell dissociation and nuclei extraction protocol respectively. Scale bars: 20um. Inset shows a representative zoomed-in cell/nucleus. **B.** Overview of experimental workflow for snRNA-seq sample submission. **C.** Representative brightfield images (40X EVOS XL) of nuclei extractions across time in differentiation. Insets depict a selected zoomed in nuclei in Figure 1A. Scale bar equals 20μm.

**Supplemental Figure 2.**
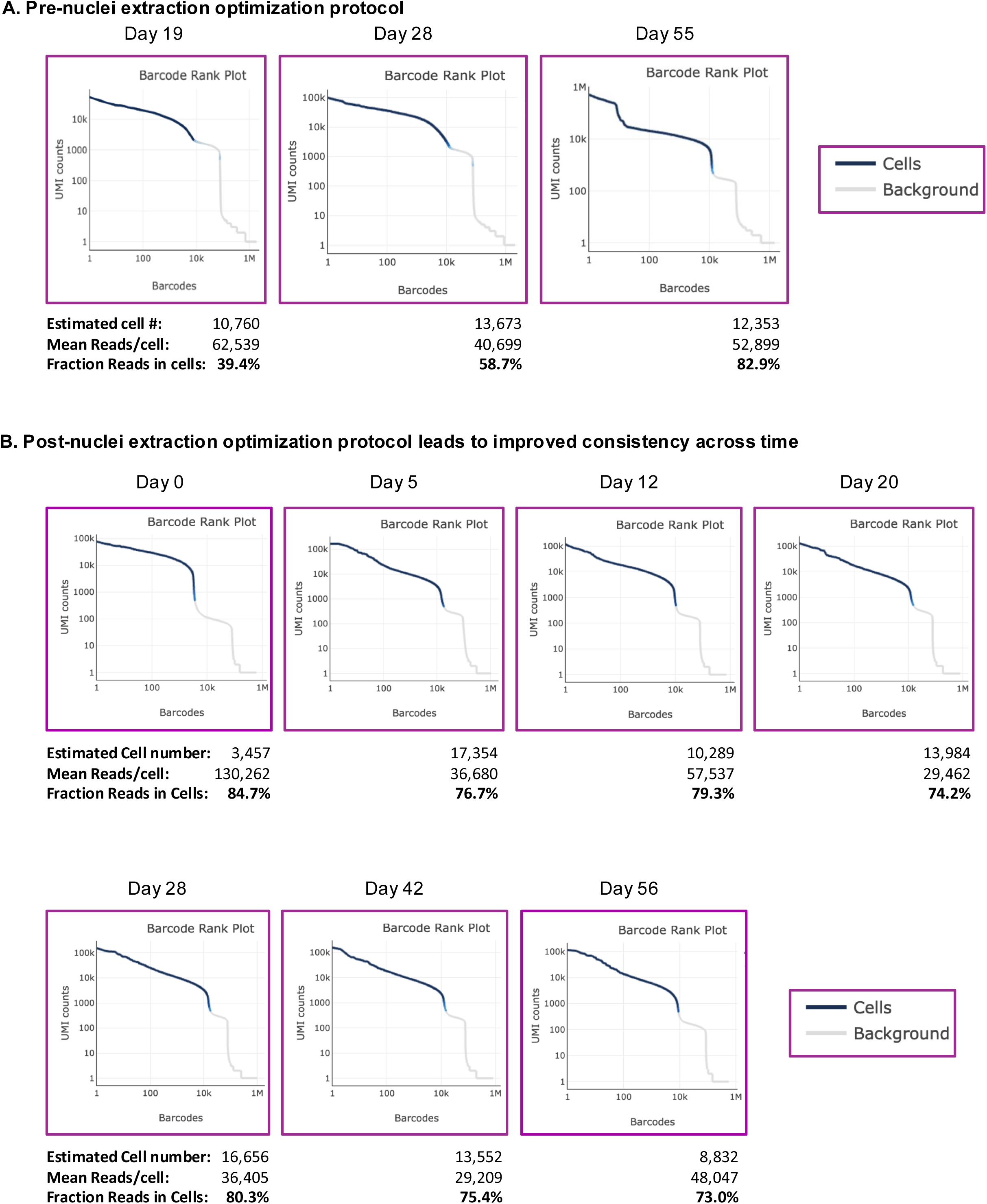
Cell Ranger plots reflect experimental quality post nuclei extraction optimizations. **A.** Cell Ranger plots before nuclei extraction protocol at days 19, 28, and 55. **B.** Cell Ranger plots after optimization of nuclei extraction protocol. Estimated cell number, mean reads per cell, and fraction reads in cells is emphasized.

**Supplemental Figure 3.**
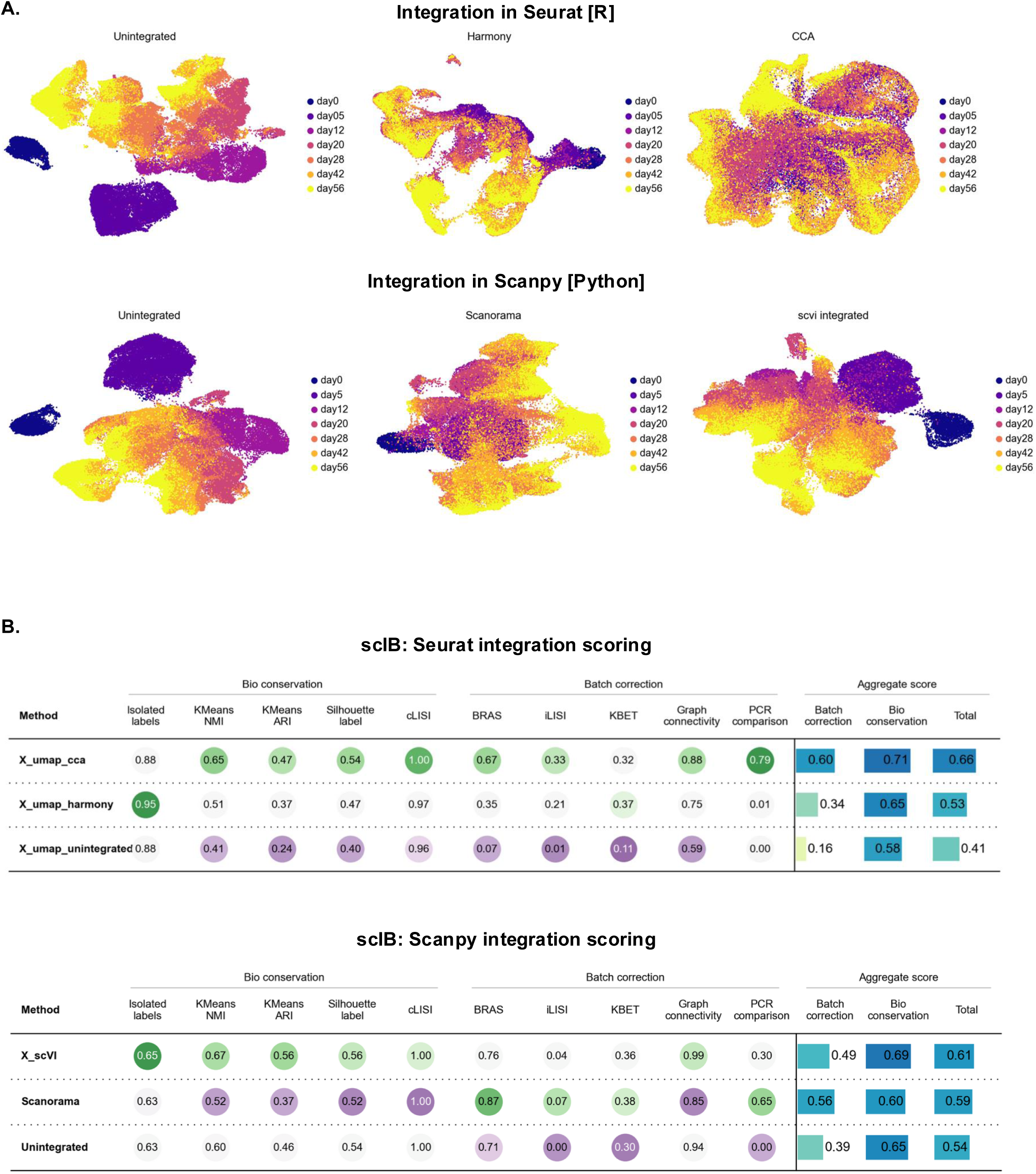
Comparison of 6 integration and unintegrated single cell methods revealed scVI performs best on complex datasets. **A.** Harmony and CCA data Integration in Seurat compared to Scanorama and scVI in Scanpy. **B.** scIB metrics to score and comparing integration methods.

**Supplemental Figure 4.**
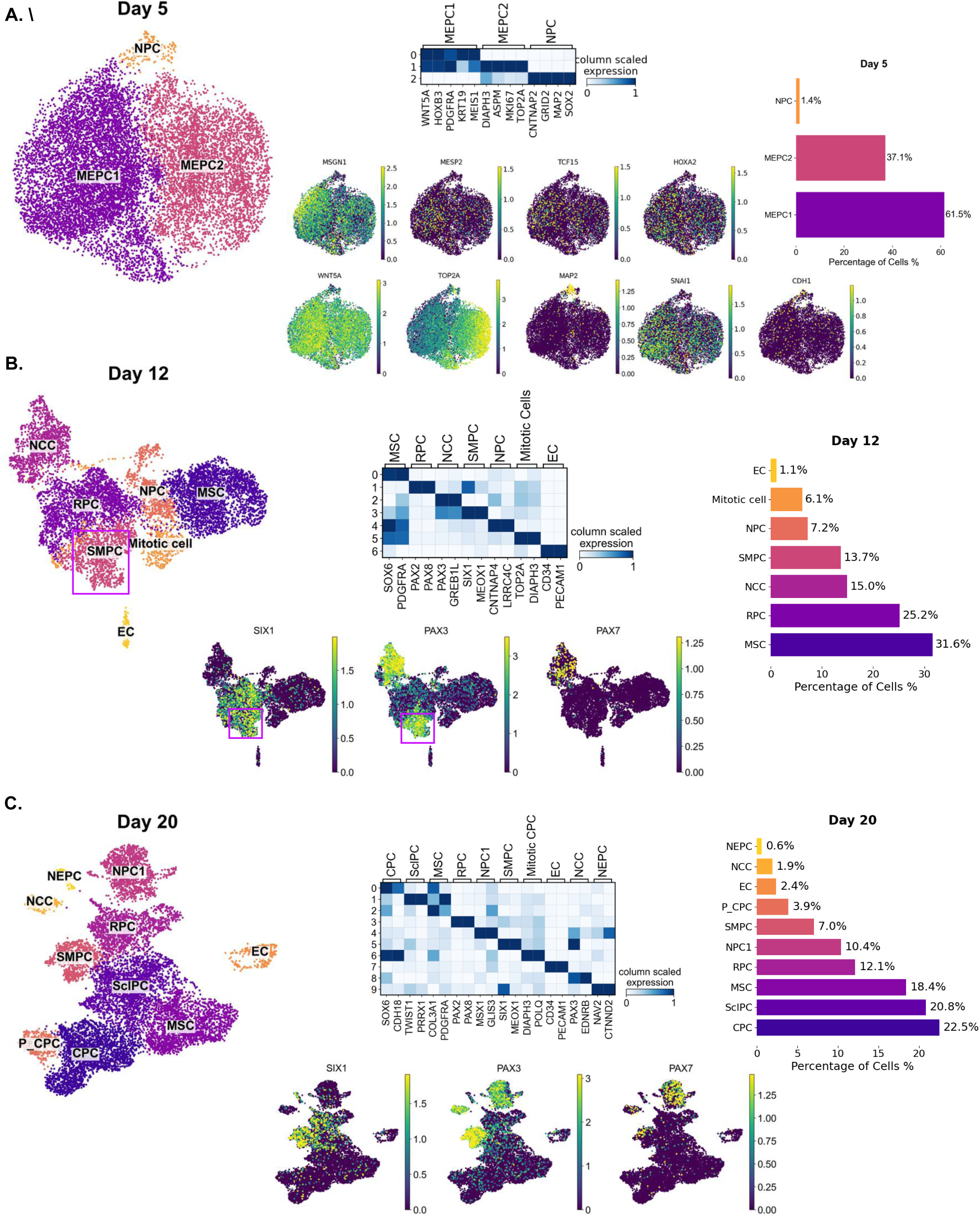
Analysis of early timepoints 5-20 refines myogenic sub-states A-C. Independent analysis of day 5, day 12, and day 20 datasets. UMAP labeled with cell type annotations and matrix plot showing the top 4 and 2 DEGS across cell types; values show the mean expression per gene grouped by cell type identity; 1 represents maximum mean expression and 0 is the minimum. Selection of key myogenic features for each time point and cell type proportions are shown. SIX1+PAX3+ SMPCs are highlighted in pink box at day 12.

**Supplemental Figure 5.**
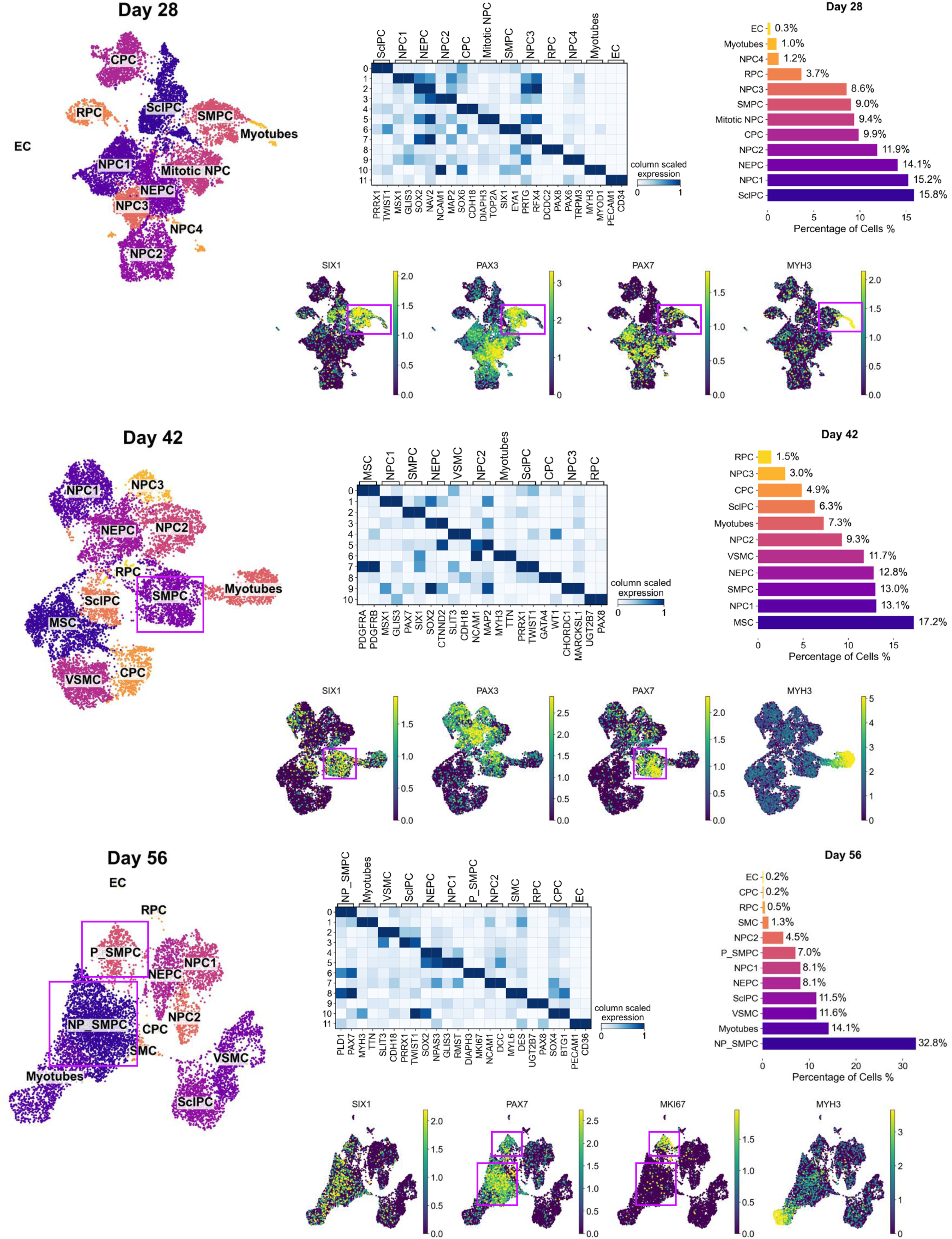
Analysis of late timepoints 28-56 refines myogenic sub-states. **A-C.** Independent analysis of days 28, 42, and 56 datasets. UMAP labeled with cell type annotations and matrix plot showing the top 4 and 2 DEGS across cell types; values show the mean expression per gene grouped by cell type identity; 1 represents maximum mean expression and 0 is the minimum. Selection of key myogenic features for each time point and cell type proportions are shown. Proliferative and non-proliferative SMPCs are defined by expression of PAX7 and MKI67 in the pink boxes.

**Supplemental Figure 6.**
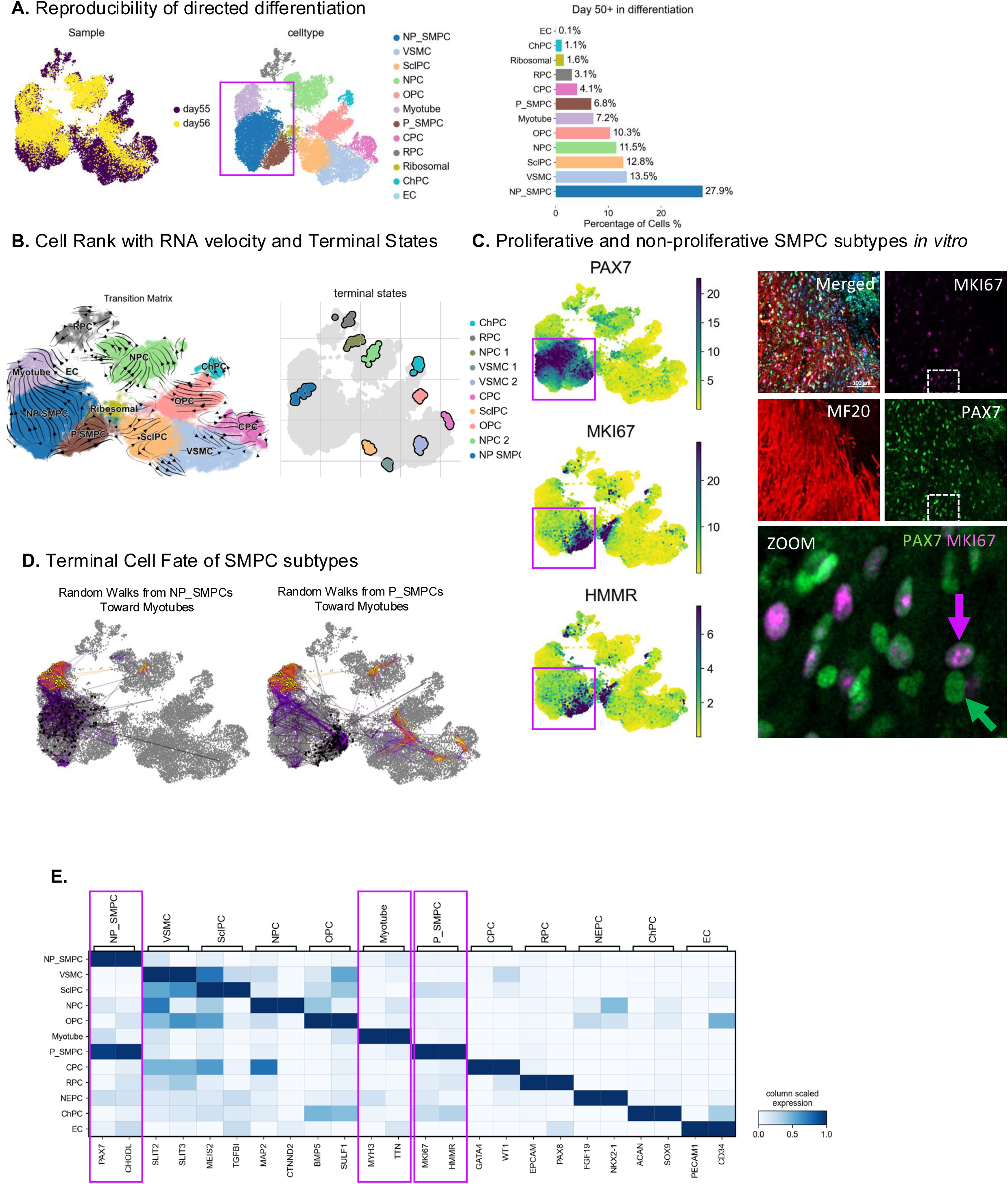
Characterization and reproducibility of terminal myogenic differentiation. **A.** *Left:* scVI normalized UMAP post integration of two independent differentiations (days 55 and 56), labeled by cell type and experimental batch. *Right:* Bar graph depicting cell type proportions (%); colors match cell type annotations. **B.** *Left*: UMAP containing scVelo projected velocities produced using CellRank’s VelocityKernel. *Right*: CellRank’s computation of terminal states using GPCCA algorithm. Cells most confidently assigned to each terminal state are highlighted. **C.** Left: PAX7, MKI67, and HMMR feature plots. *Right*: immunofluorescent images of day 50 cultures stained with DAPI (blue), PAX7 (green), MF20 (red), and MKI67 (magenta) antibodies. Scale bar: 100um. Inset shows a zoomed in image with PAX7 and MKI67 co-expression. **D.** CellRank’s random walk simulation plot showing the start (black dots) and end points (yellow) of SMPC specific lineage. *Right*: Feature plots showing expression of SMPC unique cell surface receptors. **E.** Matrix plot shows top signature DEGS for each cell type; values show the mean expression per gene grouped by cell type identity.

**Supplemental Figure 7.**
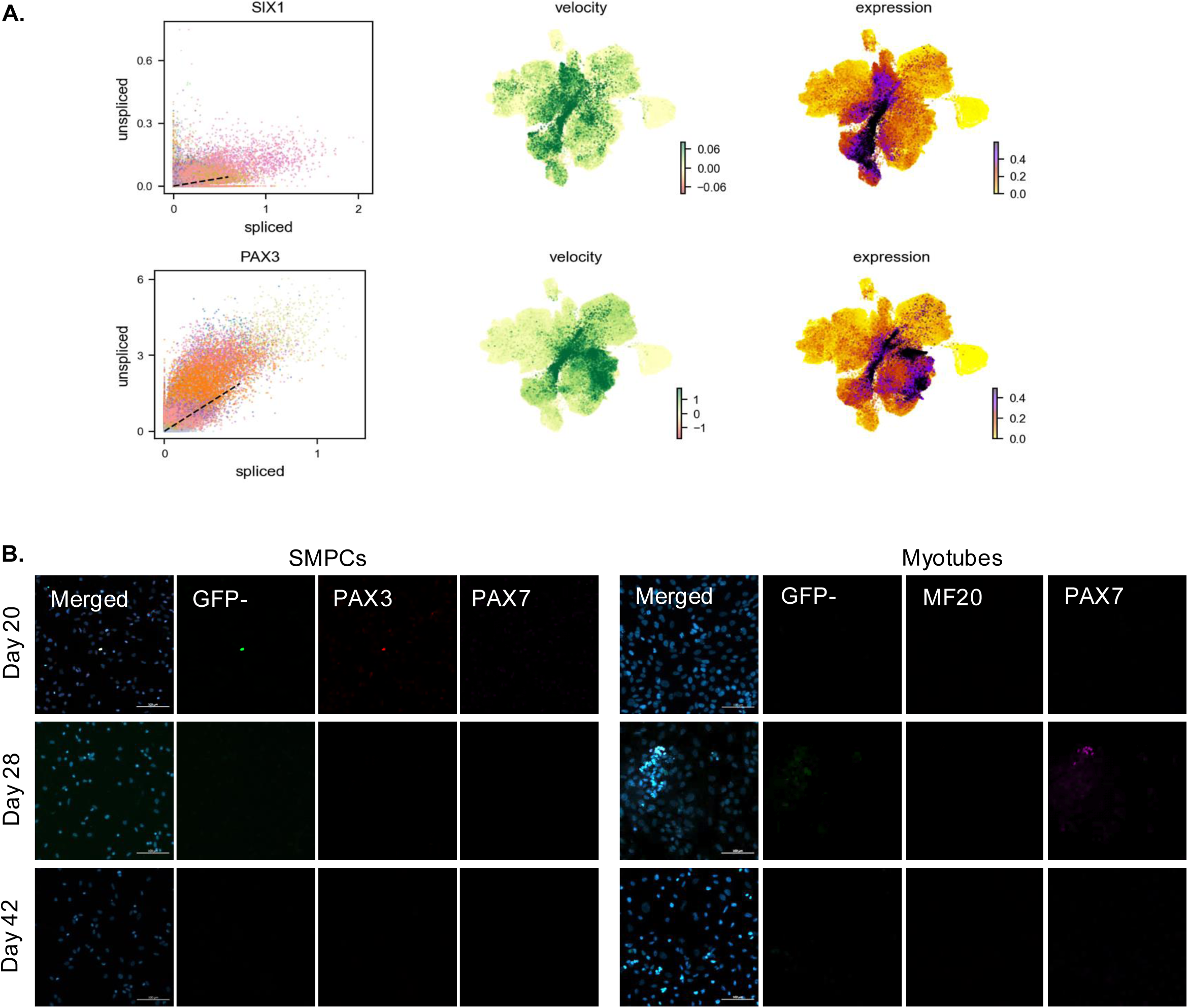
Defining SIX1 lineage trajectory for hPSC SMPCs. **A.** scVelo velocity plots for SIX1 and PAX3 measures the proportions of unspliced vs spliced RNA per cell. UMAPs showing velocity and expression patterns across time in differentiation. **B.** *Left*: Immunofluorescent images of GFP- cells stained with DAPI (blue), endogenous GFP expression (green), PAX3 (red), and PAX7 (magenta). *Right*: Immunofluorescent images of GFP- cells in myogenic differentiation media stained with DAPI (blue), endogenous GFP expression, MF20 (red), and PAX7 (magenta). Scale bars: 100um.

**Supplemental Figure 8:**
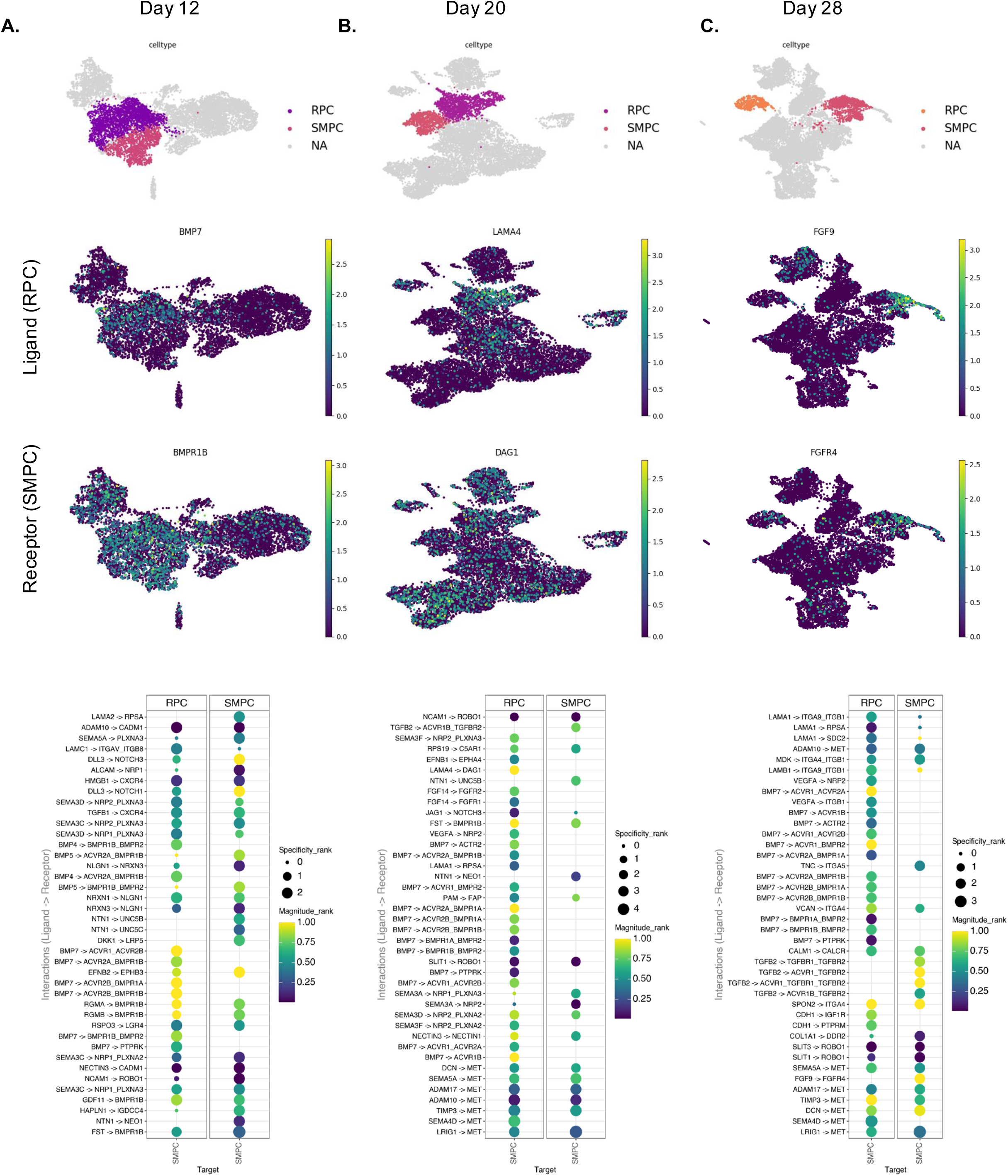
SMPCs rely on extrinsic paracrine and autocrine signals to achieve myogenic commitment. **A.** *Top:* Feature plot highlighting SMPCs and RPCs in UMAP space for day 12. Selected expression of BMP7 ligand by RPC and BMP1RB receptor by SMPC is shown. *Bottom*: LIANA+ dot plot where specificity and magnitude represent the rank of interaction specificity and strength, respectively. Larger and lighter dots correspond to specific and strong ligand-receptor interactions. **B.** *Top*: Feature plot highlighting SMPCs and RPCs in UMAP space for day 20. Selected expression of LAMA4 ligand by RPC and DAG1 receptor by SMPC is shown. *Bottom*: LIANA+ dot plot highlights LAMA4 and VEGF strong interactions. Larger and lighter dots correspond to specific and strong ligand-receptor interactions. **C.** *Top*: Feature plot highlighting SMPCs and RPCs in UMAP space for day 28. Selected expression of FGF9 ligand and FGFR4 receptors by SMPCs are shown. *Bottom*: LIANA+ dot plot highlights strong and specific SMPC autocrine signaling.

**Supplemental Figure 9:**
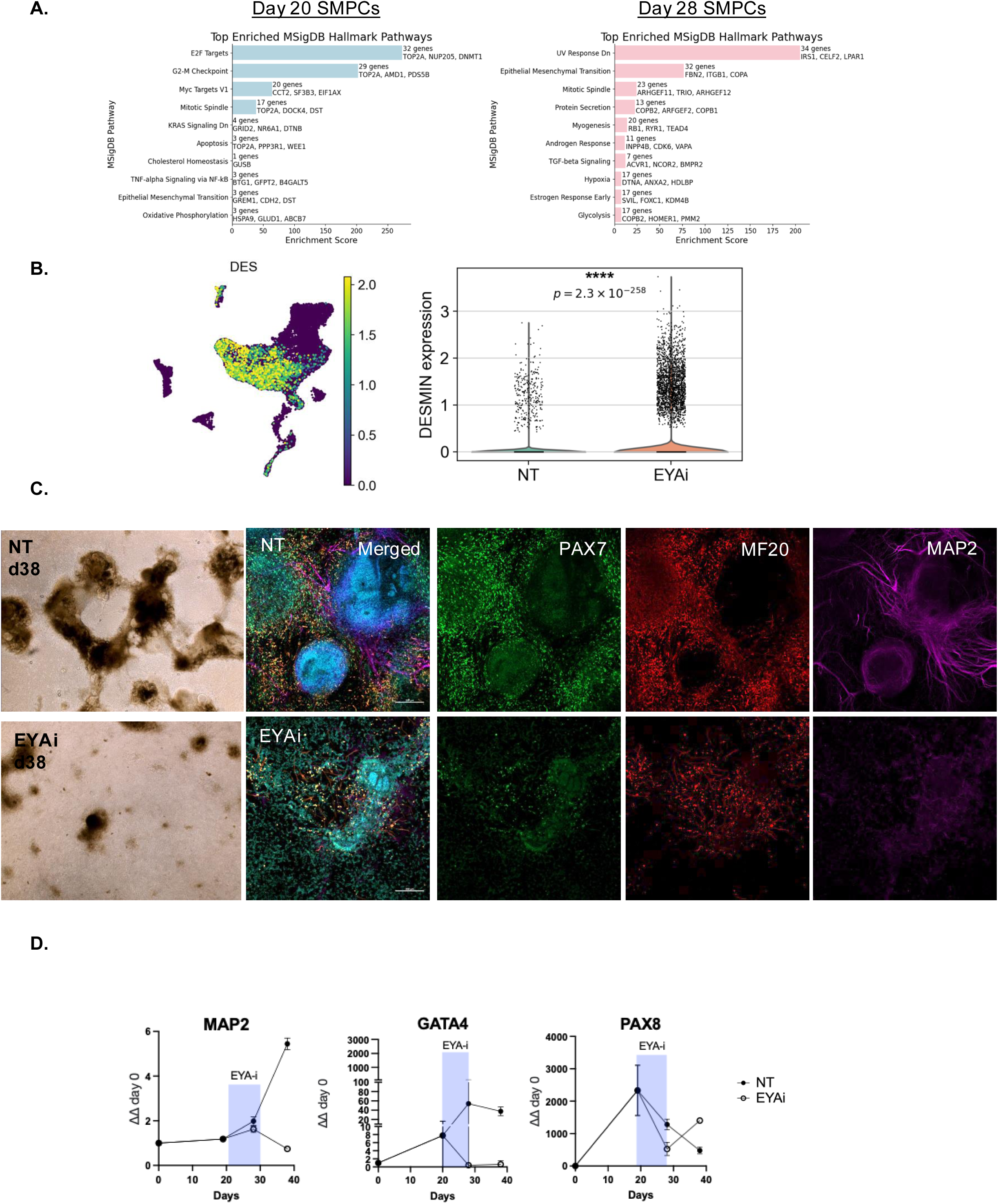
EYA inhibition leads to DES+MYH3- fibrotic cell types. **A.** Bar graph depicting EnrichR’s enriched pathways through MSigDB module; the top 3 genes and number of genes defining the pathways are highlighted from day 20 (blue) and day 28 (pink) SMPCs (see **TableS4**). **B.** Feature plot depicting expression of Desmin and violin plot showing distribution of Desmin expressing cells by treatment (P=2.3E10-258). **C.** Representative brightfield image at day 38 and IF images of non-treated (NT) and EYAi-treated directed differentiation cultures at day 28 stained with PAX7 (green), MF20 (red), and MAP2 (magenta). **D.** Quantitative RT-qPCR shows levels of MAP2, GATA4, and PAX8 in non-treated and EYAi treated culture conditions. Levels show relative expression to GAPDH and normalized to hPSCs at day 0.

## Materials and Methods

### Human Pluripotent Stem Cells (hPSC) lines and cell culture techniques

All hPSC experiments were performed with Embryonic Stem Cell Research Oversight (ESCRO) Committee approval. HPSCS were grown on Matrigel-coated plates (Corning #354277) in mTESR medium (Stemcell Technologies). HPSC lines used in this study include dCas9-KRAB (WTC11, Coriell Institute for Medical Research AICS-0090-391; Figs. 1-3, 5-7, S1-9); SIX1:H2B-GFP (H9, Memorial Sloan Kettering Institute; Figs. 4, and S6).

### Differentiation procedure

hPSCs were cultured on mTESR plus for at least three passages. Prior to the start of differentiation, hPSCs were pretreated with recovery media (10μM rock inhibitor in mTESR1 plus) for 45 min and then single cell dissociated and seeded on Matrigel coated plates at a concentration of 275,000-475,000 cells/well depending on hPSC line for differentiation. On day two, cells are treated with mesoderm differentiation media (8-10μM CHIR (Tocris Biosciences) in E6 media) for a duration of 2-3 days and then switched to SIX1+PAX3+ specification media (E6 media). Between days 5-9, cells should form 3D structures. Depending on morphology and media coloration, cells are switched to SIX1+PAX3+ expansion media (StemPro-34 + bFGF) between days 10-12 for a duration of 6-8 days. Media is then switched to PAX7 specification media (E6 media supplemented with IGF1) for 7-10 days. HPSC-SMPCs are induced to differentiate using PAX7 differentiation media (DMEM/F12 supplemented with N2 and IGF1 media for 5-7 days) and matured by addition of 3μM SB431542 to PAX7 differentiation media prior to flow cytometry. A detailed protocol on hPSC skeletal muscle differentiation including notes and tips on successfully deriving SMPCs can be found in [15].

### Nuclei Isolation for snRNA-seq

HPSCs were differentiated in a 12-well plate, with one well harvested at each of the seven timepoints for single cell dissociation followed by nuclear extraction for longitudinal tracking by single-nucleus RNA sequencing (snRNA-seq). Directed differentiation cultures at each time point were pre-treated with CEPT for 1 hr prior to single cell dissociation. Cells were dissociated by incubating with TryPLE for 5-7 min at 37°C; wide bore pipettes were used for gentle dissociation. Single cells are then centrifuged at 500g for 5 min in a chilled centrifuge (4°C). We lysed cells by incubation of ice-cold lysis buffer from the nuclei EZ prep kit (Millipore Sigma, NUC101-1KT) for 5-7 min. Lysis reaction is stopped with ice-cold 1% BSA containing RNASE inhibitor. Collect nuclei by centrifugation at 300g for 5 min. Collect single nuclei in BSA supplemented with 0.2U/μl of RNASE inhibitor. At this time, cell morphology was visualized using a 40X bright field objective on an EVOS microscope. We carried out nuclei isolations only if cells displayed healthy cell morphology (See Fig. S1A). Supernatant contains cytoplasmic components that can be saved for later analyses. Cells were resuspended in 1% BSA and passed through a 100μm filter and immediately counted using trypan blue. Single, live nuclei can further be purified using flow cytometry. Please see supplementary materials and methods for a detailed list of reagents and comprehensive protocol.

### Nuclei FACs sorting

Extracted nuclei were visualized and collected in ice-cold 1% BSA supplemented with 0.2U/μL of RNASE inhibitor and PI, then transferred to FACs tubes for FACs purification. Nuclei were sorted for single, live nuclei using the 130μm nozzle in a FACS Aria II (BD Biosciences) by the UCI Flow Cytometry Core. Sorted nuclei were visualized again for assessment of nuclei integrity and then pelleted by centrifugation for 5 min at 300g at 4 °C and resuspended at an optimal concentration of 1200 nuclei/μl in 1% BSA supplemented with RNASE inhibitor.

### snRNA-seq quality controls

10x Chromium single nuclei capture, cDNA library preparation and QC analysis (Chromium Next GEM Single Cell 3’ Kit v3.1, Cat# 1000269) was performed by the UCI Genomics Research and Technology Hub. Library preparation was performed to capture close to 10,000 barcoded cells at an average sequencing depth of 50,000 reads and 2500 genes per cell using the HiSeq 4000 Illumina platform. Cell Ranger processing of FASTQ files were mapped to the human genome GRCh38.Visualizations, clustering, and differential expression tests (Wilcoxon method) for individual time course datasets (d0, d5, d12, d28, d42, and d56) were performed using the standard Scanpy pipeline for single cell analysis in Python v3.12. (https://scanpy.readthedocs.io/en/stable/tutorials/basics/clustering.html) ENSEMBL IDs with no gene annotations were first filtered out for meaningful interpretation of the datasets. Quality control was performed by keeping number of genes by counts between >500 and <8,000, total counts <20,000 and mitochondrial percent counts <2%. For estimated number of cells and mean reads per cell captured per dataset, please see Fig. S2B.

### Data Integration of temporal datasets

Integrative analysis of the seven datasets was performed using the scVI model from scvi-tools (https://docs.scvi-tools.org/en/1.0.0/tutorials/notebooks/api_overview.html) and used for downstream trajectory analyses. Assessment of Harmony, CCA, Scanorama, and scVI integration algorithms was performed using scib-metrics in python (https://scib-metrics.readthedocs.io/en/stable/). Harmony and CCA batch correction was first performed in Seurat v5 (R v4.4.1) using the default settings and then displayed as a UMAP labeled by time. Seurat objects were then converted to anndata objects by extracting PCA, metadata, features, and matrix file and importing these into python. scVI and Scanorama embeddings were calculated in Scanpy (Python 3.12) using the recommended 3000 highly variable genes and then displayed by a batch labeled UMAP. All four integration methods were then benchmarked using Leiden/Seurat-clusters through scIB algorithm. Aggregate scores taking into consideration both conservation of the biological data and batch correction scores were used to evaluate how well clusters matched the underlying data structure and how well cells of the same cell type were connected.

### Cell-type annotation

Prior to cell annotation, we re-assessed quality control and cell filtering. We manually annotate cell types using genes exclusively expressed by a given cell cluster. We focus on marker genes distinguishing a cluster from the heterogeneous groups of cells by using Scanpy’s ‘rank_genes_groups’ function using Wilcoxon method. By plotting the top 25 genes, we visualize clear patterns of expression. We further obtained cluster-specific differentially expressed genes and use these genes to manually determine cell type. We use a combination of tools and available resources to compare these genes across existing datasets (EnrichR, PanglaoDB, CellMarker). Using a predefined gene list, EnrichR generates a list of enriched terms through its comprehensive and diverse gene sets where each library is composed of a group of related gene lists associated with a pathway, ontology, disease, cell type, or transcription factor. We also relied on our background knowledge (myogenic cells), and published literature on known definitive marker genes.

### Percent contribution of cells expressing a gene

Percent expression was defined as the fraction of cells with at least 1 detected UMI for a given gene. Computation of percent of cells expressing a specific gene was calculated from raw count data prior to normalization or feature selection.

### Moscot trajectory analysis

To obtain a more accurate mapping of the myogenic lineage trajectory, the scVI-integrated dataset was filtered to the day 0 and day 5 progenitors, and myogenic cell types across time. The TemporalProblem tutorial was followed to compute the cost matrix (Epsilon=1e-3, max iterations=1e7) to generate transition matrices and visualizations of ancestors and descendants. Identification of driver genes and transcription factors (features=human) was performed by computation of the pull distributions at each time point. https://moscot.readthedocs.io/en/latest/notebooks/tutorials/200_temporal_problem.html

### scVelo/scVI

RNA velocity analysis was performed using scVelo (v0.2.3) in Python. We first prepare the scVI integrated dataset by loading the loom files (generated with Cell Ranger) with its corresponding dataset to obtain splicing information matrices (ldata0 = sc.read(‘loom_file_directory’, cache=True). Then we run scVelo’s (v0.2.3) dynamical model for downstream analysis with CellRank. https://scvelo.readthedocs.io/en/stable/DynamicalModeling.html

SCVI model ran with the following parameters: n_hidden: 128, n_latent: 10, n_layers: 1, dropout_rate: 0.1, dispersion: gene, gene_likelihood: zinb, latent_distribution: normal.

Training status: Not Trained; Model’s adata is minified? False. ScVI embedding was normalized using scanpy (leiden resolution = 0.6).

### CellRank

The CellRank Meets RNA velocity tutorial was followed to generate velocity projected UMAPs and random walk plots. We first run scVelo’s dynamical model and then set up the velocity kernel to compute a transition matrix using the deterministic mode. The velocity kernel is then combined gene expression similarity to generate a connectivity kernel. We visualize transition matrix using UMAP and generate random plots (max iterations=200, seed=0) to observe transitions of specific cells. Black dots define starting points, and yellow dots represent ending points to qualitatively visualize transition paths. https://cellrank.readthedocs.io/en/latest/notebooks/tutorials/kernels/200_rna_velocity.html The velocity kernel was used to then compute initial and terminal states using the GPCCA method. https://cellrank.readthedocs.io/en/latest/notebooks/tutorials/estimators/600_initial_terminal.html

### Gene enrichment and pathway analyses

Differential gene expression analysis was first calculated in Python using the Scanpy single-cell analysis framework. We identified differentially expressed genes (DEGs) between specified cell populations using Scanpy’s built-in testing functions with multiple-testing correction (Benjamini–Hochberg), and genes with an adjusted p-value (P_adj) < 0.05. DEG lists were then uploaded to EnrichR, https://maayanlab.cloud/Enrichr/. For pathway-level interpretation, we queried the Molecular Signatures Database (MSigDB) collections to identify enriched biological processes and signaling pathways and used PanglaoDB gene sets to relate DEGs to cell type–specific marker signatures. Enrichment was assessed using Enrichr’s default statistical settings, and results were summarized based on combined scores and adjusted p-values.

### LIANA+

Steady state ligand-receptor inference was performed using LIANA+ tutorials: https://liana-py.readthedocs.io/en/latest/notebooks/basic_usage.html. Briefly, the processed single-cell RNA-seq object at each timepoint (AnnData) with cell-type annotations was used as input for LIANA pipeline. We applied multiple underlying methods implemented in LIANA+ (e.g., CellPhoneDB, Connectome, NATMI) and combined their outputs using the rank aggregate (RRA) approach to derive a consensus ranking of ligand-receptor pairs across methods. Interactions were filtered to retain those with adequate expression in both sender and receiver cell types according to LIANA+ default thresholds and ranked by the aggregated “magnitude rank” score, which reflects the relative strength and consistency of each interaction. For visualization, dot plots display the top 10 or 40 most relevant ligand–receptor interactions, ordered by aggregated magnitude rank, with dot size and color representing interaction strength and/or expression metrics as indicated in figure legends.

Code availability and details can be found on GitHub: https://github.com/mhickslab/Reconstruction-of-hPSC-Myogenic-Lineage-Transitions-through-Single-Cell-Optimal-Transport-

### Microscopy

Images were taken on a Zeiss LSM900 Airyscan confocal microscope. 6-well/12-well plastic plates were imaged using the 10X objective with either 0.5x or 1.5x zoom. Images were taken with a maximum laser intensity of 5-10% and voltage 600-800V. Maximum intensity projection and airyscan processing were applied.

### Fluorescence Activated Cell Sorting

Cells were single cell dissociated using wide bore pipettes in TrypLE (Gibco with no phenol red), neutralized with CEPT supplemented media (E6, N2, or SKGM2), and filtered through 100µm filters. Single cell suspensions were centrifuged at 500g for 5 minutes. Pellets were resuspended in FACS buffer (2% FBS in dPBS) and stained with viability dye (PI) at a 1:1000 dilution. Samples were transferred through 70µm 5mL FACS tubes and immediately sorted for GFP+/− populations using the FACS ARIA FUSION sorter.

### EYA3 inhibition Assay

dCas9-KRAB hPSCs were differentiated to skeletal muscle under normal culture conditions. following our established protocol. Between days 20-28, treatment with either 6-benzbromarone at 3uM for EYAi group or DMSO vehicle control for NT group was added every day with their corresponding differentiation media. At day 28 of differentiation, the treatment ended and all groups of cells (NT, EYAi) were continued to be grown under normal conditions until day 38 of differentiation.

### Immunocytochemistry

All cells were fixed with 4% paraformaldehyde (PFA) for 15 mins and permeabilized using 0.5% Triton X-100 for 15 mins. Fixed cells were blocked in 5% bovine serum albumin with 10% fresh goat serum for 1 h. Cells were incubated overnight at 4°C in primary antibody solution containing fresh 10% goat serum. Following incubation, cultures were washed three times using PBS, followed by secondary antibody incubation containing fresh 10% goat serum for 1 h at room temperature. See supplementary materials and methods for vendor and dilution information.

### RT-qPCR

Validation of RT-qPCR primers was conducted by performing primer efficiency curves. When collecting cell pellets for RT-qPCR, it is necessary to filter excess ECM contents using 70-100 µm filters from SMPC-derived cultures, as these tend to clog RNAeasy columns and affect RNA yield quality. Total RNA was collected using Qiagen RNeasy Plus Mini Kits with an adjusted protocol and RNA yield and quality was analyzed using a NanoDrop 2000 spectrophotometer. cDNA was synthesized using iScript Reverse Transcription Supermix (Bio-Rad), and gene expression was determined using a Quantstudio 6 Flex Real-Time PCR System with SYBR Green PCR Master Mix (Bio-Rad). Experiments were conducted using 384-well plates, with the amounts of SYBR Green, RNAse free water and primers adjusted accordingly. RT-qPCR data was analyzed using ΔΔCT in which every hPSC line compared with its own day 0 pluripotent control. See Supplementary Materials and Methods for detailed protocol and primer sequences.

